# Integrative solution structure of a PTBP1-viral IRES complex reveals strong compaction and ordering with residual conformational flexibility

**DOI:** 10.1101/2022.07.08.498958

**Authors:** Georg Dorn, Christoph Gmeiner, Tebbe de Vries, Emil Dedic, Mihajlo Novakovic, Fred F. Damberger, Christophe Maris, Esteban Finol, Chris P. Sarnowski, Joachim Kohlbrecher, Timothy J. Welsh, Sreenath Bolisetty, Raffaele Mezzenga, Ruedi Aebersold, Alexander Leitner, Maxim Yulikov, Gunnar Jeschke, Frédéric H.-T. Allain

## Abstract

RNA-binding proteins (RBPs) are crucial regulators of gene expression and often comprise well-defined domains interspersed by flexible, intrinsically disordered regions. The structure determination of ribonucleoprotein complexes involving such RBPs is not common practice and requires integrative structural modeling approaches due to the fact that they often do not form a single stable globular state. Here, we integrate data from magnetic resonance, mass spectrometry, and small angle scattering to determine the solution structure of the polypyrimidine-tract binding protein 1 (PTBP1 also called hnRNP I) bound to an RNA which is part of the internal ribosome entry site (IRES) of the encephalomyocarditis virus (EMCV). PTBP1 binding to this IRES element enhances translation of the viral RNA. The determined structural ensemble reveals that both RNA and protein experience a strong compaction upon complex formation, get ordered but still maintain a substantial conformational flexibility. The C-terminal RNA recognition motif (RRM4) of PTBP1 rigidifies the complex by binding a single-strand RNA linker and, in turn, is essential for IRES-mediated translation. PTBP1 acts as an RNA chaperone for the IRES, by ordering the RNA into a few discrete conformations that expose the RNA stems to the outer surface of the RNP complex for subsequent interactions with the translation machinery. The conformational diversity within this structural ensemble is likely common among RNP complexes and important for their functionality. The presented approach opens the possibility to characterize heterogeneous RNP structures at atomic level.

## Introduction

Gene expression is critically regulated by protein-RNA interactions. RNA-binding proteins (RBPs) usually contain multiple RNA-binding domains (RBDs), among which the RNA recognition motif (RRM) is the most abundant domain type (1). Typically, RBDs are flanked and linked by intrinsically disordered regions (IDR) of various length and sequence (2). Recently, these flanking IDRs have been studied extensively in the context of liquid-liquid phase separation (3-5). For example, the low complexity domain of the ALS-associated protein FUS has not only been connected to functional phase separated states, but also associated with plaque formation that causes neurotoxicity (6, 7). The IDRs that link RBDs potentially have diverse biological function and they unavoidably cause flexibility and heterogeneity of the overall protein conformation. However, cooperative binding of the RBDs to RNA binding sites may result in an increased conformational order of the protein. This evokes the questions to which degree a disorder-to-order transition is achieved, and if both flexibility and rigidity are important for the biological function of the protein-RNA complex.

A plethora of structures of isolated RBDs are available from nuclear magnetic resonance (NMR) and X-ray crystallography (see examples in refs (8-14)). They generally represent well-defined, three-dimensional folds following the Anfinsen dogma (15). However, the presence of structurally undefined flexible regions connecting these subunits and the associated conformational heterogeneity hinders the structure determination of full-length RBP and their complexes by using exclusively these classical techniques, as well as cryo-EM (16-18). Thus, current access to ribonucleoprotein (RNP) structures is limited to protein-RNA machineries with stable intermediates such as ribosomes and spliceosomal particles or to protein-RNA complexes of mainly single and tandem RBDs bound to optimized RNA sequences (19-21). Yet, the biological function of protein-RNA complexes may depend on adoption of multiple conformations, which may be important for binding distinct targets (14, 22). Integrative structural biology approaches combining multiple methods overcome the restrictions of classical methods in characterization of such heterogeneous structures (18). However, the integration of a variety of structural constraints with different precision and length scales, as well as, the aforementioned features of the protein-RNA complexes also require a change in the structure determination concept at the modeling stage.

The abundant 57 kDa RBP polypyrimidine tract protein 1 (PTBP1, also hnRNP I) is a general regulator of cellular mRNA metabolism. In the context of splicing regulation, PTBP1 acts either as splicing repressor, potentially by binding competition with other factors, or as splicing activator in a position-specific manner (23-27). PTBP1 also determines the localization, stabilization and polyadenylation of many target RNAs and is involved in the translational regulation of cellular and viral mRNAs as an prototypical internal ribosome entry site (IRES) trans-acting factor (ITAF) (28). IRES elements allow non-canonical translation initiation in a 5’-cap independent manner. These large, highly structured RNA sequences are present in the 5’ untranslated regions (UTRs) of particular cellular mRNAs and of the genomes of positive-strand RNA viruses. IREScs enable the translation during global repression of canonical translation due to cell stress. For instance, during viral infections canonical translation is arrested and non-canonical translation activates the antiviral response and programmed cell death to prevents further viral spread. In this context, PTBP1 acts as an ITAF and favors the translation of important pro-apoptotic factors (29-31). Similarly, RNA viruses have evolved PTBP1-responsive IRES elements in their 5’ UTRs to hijack the antiviral response and favor the translation of their viral proteins (32, 33). PTBP1 is among the most frequently found ITAFs and it is an extensively characterized enhancer of the IRES-mediated translation in picornaviruses, including poliovirus (PV), human rhino virus (HRV), hepatitis C virus (HCV), foot and mouth disease virus (FMDV) and several cardioviruses such Theiler’s murine encephalomyelitis virus (TMEV) and encephalomyocarditis virus (EMCV) (34). Structurally, PTBP1 consists of four RRMs, a flexible N-terminus, which contains both nuclear localization and nuclear export signals, and IDR linkers connecting RRM1 with RRM2, as well as, RRM2 with RRM3, respectively (8, 35, 36) (Fig. 1A). The nuclear magnetic resonance (NMR) solution structures showed that each of the four RRMs adopts the classical RRM topology of βαββαβ, which is extended by a fifth β-strand in RRM2 and RRM3, and that each domain binds to specific cytosine/uridine-rich sequences within single-stranded RNA (10, 37). It has been proposed that PTBP1 could act as an RNA chaperone since RRM3 and RRM4 stably interact, thereby spatially restricting the orientation of the cognate RNA binding sites (8, 35). Recent NMR studies of the individual RRM1 and RRM2 bound to UCUUU pentaloops illustrated the capability of PTBP1 to bind to structured RNA targets and identified an α-helix at the C-terminus of RRM1 as of importance for sensing RNA secondary structure (38, 39). However, the interplay of the four domains and their assembly on a natural RNA target remain unknown. Understanding the RNA-binding mode of PTBP1 is important, as misregulation of its expression has been involved in disease promotion, including colorectal cancer invasion, breast cancer cell growth and Parkinson’s disease (40-42).

**Fig. 1.**
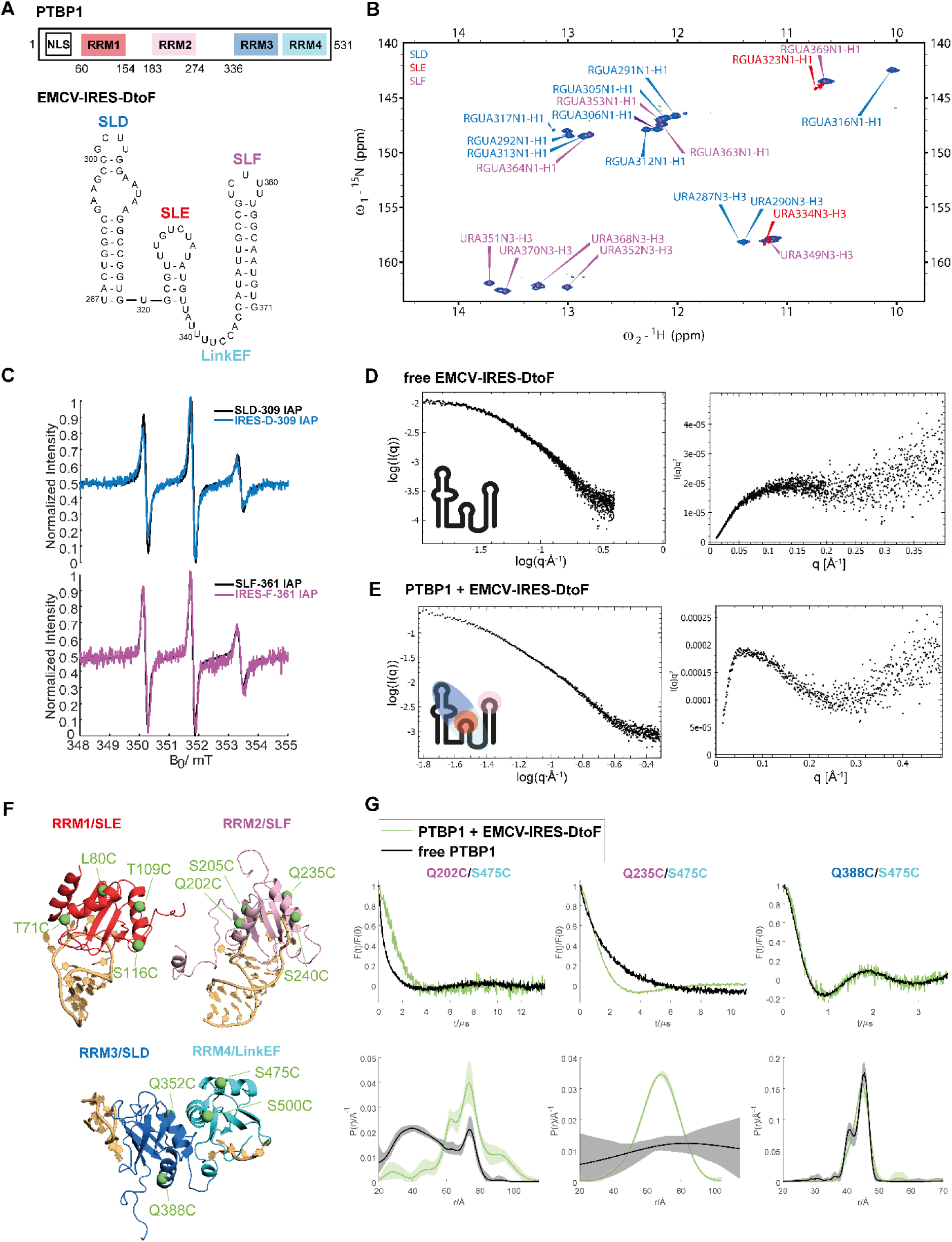
PTBP1 and EMCV-IRES-DtoF are flexible in the free state and rigidify within the complex. (A) Domain scheme of PTBP1 and overview of the EMCV-IRES-DtoF RNA construct used in this study. (B) 2D ^1^H-^15^N TROSY spectrum of EMCV-IRES-DtoF imino cross-peak signals confirm base-pairing. Imino cross-peaks of G323 and U334 of SLE shifted as indicated by red arrows. (C) CW-EPR spectra of SLD and SLF (black) and EMCV-IRES-DtoF (magenta) with spin label attached to nucleotide 309 and 361, respectively. The sharpness of the signals does not change, implying a similar flexibility of the label in the single stem loop RNA as in the whole EMCV-IRES-DtoF. (D) The SAXS data measured on free EMCV-IRES-DtoF shows in the Kratky plot (right) characteristics of a flexible and unstructured molecule. (E) The SAXS curve of the PTBP1-EMCV-IRES-DtoF complex shows in the Kratky plot (right) characteristics of an overall structured molecule with reduced flexibility compared to the free RNA shown in (D). (F) Overview of spin label attachment sites on the individual RRMs of PTBP1 used in this study. Additionally, spin labeling sites were placed in the linker region between RRM1 and RRM2 (152 and 156) and RRM2 and RRM34 (288, 315, 327), respectively. (G) Selected DEER-EPR measurements using spin labels attached to RRM2 (202, 235), RRM3 (388) and RRM4 (475). In the free state (black) signals decay smoothly (top panel) reflecting a broad distance distribution (bottom panel) with the exception of RRM3-RRM4, which interact stably and thus, lead to an oscillatory decay corresponding to a narrow distance distribution. Upon complex formation (green), the mean distance between RRM2 and RRM4 can increase (202/475) or decrease (235/475), while there is no significant change upon binding for RRM3/RRM4 (388/475). These distance distributions reflect a more ordered and rigidified arrangement of the RNA within the complex.

Previous studies of PTBP1 in complex with the IRES of EMCV (834 nucleotides, nts) suggested a protein-RNA binding ratio of 2:1 with binding of one molecule of PTBP1 to the IRES stem-loops (SLs) D-F (herein referred to as EMCV-IRES-DtoF, Fig. 1A) and of a second molecule to SLs H-L (43). Mutation of the RNA interface of the individual RRMs reduced IRES activity for RRM1, RRM3 and RRM4 and abrogated IRES activity for RRM2 and RRM34 (44). Interestingly, mutation of the RNA interface of an individual RRM did not change the binding of the other RRMs. Recently, we could revise the interaction pattern of PTBP1 with EMCV-IRES-DtoF and proposed three-dimensional models of the individual PTBP1 RRMs in complex with parts of the RNA (39).

Here, we applied an integrative structural biology approach combining electron paramagnetic resonance (EPR), NMR, small angle neutron and X-ray scattering (SANS/SAXS), and cross-linking of segmentally isotope labeled RNA with tandem mass spectrometry (CLIR-MS/MS), to study the structure of the full-length PTBP1 in complex with an RNA derived from the IRES of EMCV (EMCV-IRES-DtoF, 84 nts). PTBP1 and EMCV-IRES-DtoF are highly dynamic molecules in their isolated states, and assemble together into a compact and ordered structure, yet still sampling a fairly large conformational space. This conformational variety can only be described by a structural ensemble. As confirmed using EPR and bicistronic reporter assays, binding of RRM4 to the single-stranded RNA-linker between SLE and SLF is crucial for a stable complex assembly as well as for IRES activity, consistent with an RNA chaperone role of PTBP1 in enhancing IRES-mediated translation.

## Results

### PTBP1 and the EMCV-IRES RNA are flexible molecules in solution

As a first step towards the structural characterization of PTBP1 in complex with EMCV-IRES-DtoF, we analyzed the RNA sequence conservation within cardioviruses and the structural conformations of the protein and the RNA in their isolated states using NMR, EPR and SANS/SAXS (Fig. S1). The bioinformatics analysis of the EMCV-IRES-DtoF RNA (Fig. S1C,D) suggested an evolutionary conservation of the stem-loops D and F (herein referred to as SLD and SLF, Fig. 1A), while the stem of stem-loop E (SLE) appeared to be preserved only in EMCV-related viruses. The PTBP1 binding sites for all four domains (39) in the RNA (pyrimidine-tracts) are also conserved in cardioviruses (Fig. S1C,D). Using NMR, the predicted secondary structure of EMCV-IRES-DtoF were validated through the detection of imino proton signals, which report on the base-pairing within the RNA SLs. All expected imino protons of stem-loops D, E and F were assigned by assignment transfer from the individual SLs to EMCV-IRES-DtoF (Fig. 1B, Fig. S1A,B). The imino proton signals of the individual SLD (except G317) and SLF overlapped perfectly with their corresponding signals in the EMCV-IRES-DtoF RNA indicating the same fold. Imino proton resonances for the G323-U334 base-pair of SLE were detectable, even though the U334 signal was slightly shifted compared to the isolated SLE that embedded a longer stem (Fig. 1B, Fig. S1B). This indicated that the four base-pairs containing stem (321-324/333-336) exists in the EMCV-IRES-DtoF construct, and that, overall, the predicted RNA secondary structure forms in solution, in agreement with its evolutionarily conservation (Fig. S1C,D). By continuous wave (CW) EPR experiments, we observed that paramagnetic spin labels (iodoacetamido proxyl, IAP) attached to isolated SLs reveal the same EPR line shapes compared to corresponding positions within the entire EMCV-IRES-DtoF, reflecting that the tumbling rate of the spin labels between individual SLs and the full-length RNA construct remained unaffected (Fig. 1C). The small peak-to-peak line widths in these spectra, as well as the nearly symmetric EPR lines showed that spin labels are in the fast tumbling regime when they are attached to RNA. We thus concluded that in its free state, EMCV-IRES-DtoF is a highly flexible molecule. This conclusion was supported by SAXS measurements recorded on the free EMCV-IRES-DtoF, which resulted in a Kratky plot showing a plateau and no maximum, which is characteristic for highly flexible molecules (45) (Fig. 1D). Taken together, NMR, EPR and SAXS data show that the RNA in its free form adopts the predicted secondary structure with a large degree of overall conformational flexibility.

Acquiring distance information by EPR requires site-directed spin labeling at two sites within the protein or RNA. For proteins, we exploited methanethiosulfonate spin labeling (MTSSL) or IAP spin labeling and used the four-pulse double electron-electron resonance pulse (DEER) experiment for measuring electron spin-spin distance distributions (46, 47). The inter-domain DEER measurements between spin-labeled sites located in RRM2 and RRM34 revealed broad distance distributions (Fig. 1F,G). This implies the absence of any defined placement of RRM2 with respect to RRM34, consistent with an independent tumbling of RRM2 (as well as RRM1) as noted in previous NMR studies (8). In contrast, the distance distribution between spin labels attached to RRM3 and RRM4 reflected the well-defined mutual arrangement of the two RRMs by hydrophobic interactions between the α-helices of the RRMs as described earlier (35). The CW EPR spectra of MTSSL attached to PTBP1 revealed a much slower rotational tumbling regime. This is consistent with the overall tumbling of rigid RRMs, which are bulkier than RNA SLs. Analysis of the small angle neutron scattering (SANS) curves suggested an elongated shape for the full-length PTBP1, as already described earlier by others (48).

In summary, the NMR, EPR and SANS/SAXS data show that the free forms of both the RNA and protein are best described by rigid, well defined bodies (RRMs or SLs) connected by flexible linkers that allow fluctuating spatial arrangements.

### PTBP1 and EMCV-IRES-DtoF form a compact complex with residual dynamics

Complex formation between PTBP1 and EMCV-IRES-DtoF causes drastic changes in the structural arrangement of both components. Inter-domain distance distribution measurements by DEER showed clear oscillations in the dipolar evolution functions, corresponding to significantly narrower distance distributions compared to the respective free states (Fig. 1G). This clearly indicated a preferred domain-to-domain orientation within the complex. Considering that all domains bind distinct sites on the RNA (39), this implies that the RNA structure is also stabilized in a preferred conformation. Importantly, the local secondary structure of the RNA remained unchanged upon PTBP1 binding since the imino proton signals are identical compared to the free state (Fig. S1E). In total, we measured a set of 35 DEER distance distributions between 17 spin labeling sites on doubly-labeled PTBP1 in complex with native EMCV-IRES-DtoF (Fig. S2, Table S2). Interestingly, not all selected spin label pairs in PTBP1 result in narrow distance distributions in the complex, and some spin pairs exhibit standard deviations of distance of up to 28.1 Å. This exceeds the combined conformational flexibility of the two labels by far and thus reflects a preserved flexibility of the complex. Therefore, although PTBP1/EMCV-IRES-DtoF complex undergoes rigidification and compaction, it also retains certain dynamics and is likely to adopt multiple conformations. The disorder-to-order transition upon binding is incomplete. Interestingly, DEER measurements between residues located in the RRMs and the protein linkers reveal that the distance distributions between RRM1 or RRM2, and their connecting linker (Link12) are narrower than between RRM2 or RRM3, and Link23 (Fig. S2, Table S2). Hence, Link12 seems to be more restrained in the complex state and less flexible than Link23. The results from EPR and NMR spectroscopy are in line with SAXS data on PTBP1/EMCV-IRES-DtoF and the free RNA. The Kratky plot of the complex compared to the free RNA (Fig. 1E) reflects a more structured molecule (45), confirming that the IRES RNA, which contributes dominantly to the scattering, is rearranged upon PTBP1 binding.

### Structure determination of the PTBP1-EMCV-IRES-DtoF complex

We developed a structure determination protocol to incorporate both short-range distance restraints from NMR and CLIR-MS/MS and inter-domain, long-range distances from EPR and SANS/SAXS to calculate the structure of the PTBP1/EMCV-IRES-DtoF complex. The approach was built on our previously introduced methodology (49, 50). Here, we essentially defined rigid bodies comprising RRM/RNA sub-complexes, which were positioned with respect to each other by EPR long-range distance restraints using the simulated annealing structure calculation algorithm CYANA (51). The structures of conformers in the initial ensemble were refined by YASARA and the whole ensemble was fit against EPR and SAS data (Fig. S3).

The following observations allowed us to confidently define that the PTBP1/EMCV-IRES-DtoF complex consists of three rigid building blocks connected by flexible peptide and RNA linkers. First, through a combination of NMR and CLIR-MS/MS data for each of the four RRMs of PTBP1, we could place their location to a unique RNA part of the EMCV-IRES-DtoF (39). The NMR spectra of the full-length PTBP1 bound to EMCV-IRES-DtoF overlap with those of the individual domains bound to their respective RNA targets, suggesting no apparent interactions between the individual domains themselves and with the inter-domain linkers or the N-terminus. This analysis also confirmed that the structures of individual RRMs bound to their RNA targets are preserved in the full-length protein. Second, as a basis for performing site-directed spin labelling (SDSL) and EPR-based distance measurements on the full-length PTBP1 protein, we demonstrated that the EPR-based distance distributions within each isolated RRM are consistent with the NMR-based structures (47). Importantly, the distance distributions between the labeled positions within each RRM of the full-length protein were identical to those measured on the isolated RRMs (Fig. S2). Finally, as shown above, complex formation does not alter the secondary structure of the EMCV-IRES-DtoF RNA (Fig. S1E).

Accordingly, we defined RRM1 bound to SLE, RRM2 bound to SLF and RRM34 bound to SLD (RRM3) and to four nucleotides of the LinkEF (RRM4) as rigid bodies. The structural details of these sub-complexes have been determined previously using a hybrid approach of NMR, CLIR-MS/MS and available structural data (39). Uncertainties in the RRM1/SLE model were resolved by additional NMR analysis, including the assignment of intermolecular nuclear Overhauser effects (NOEs) to confirm the binding register (Fig. S4). Structures of the individual RRMs and SLs were used as input for the structure calculation in CYANA and kept rigid by fixing the backbone torsions within the corresponding segments throughout the calculation. To form RNA stems, we used hydrogen-bond restraints while loops and bulges were maintained flexible. Intermolecular restraints between RRMs and SLs were incorporated to obtain arrangements as in the previously determined structures. In addition to the 35 long-range, protein-protein EPR distance distributions between these three individual sub-complexes, we additionally recorded SAXS data as well as a SANS contrast variation series. The EPR-derived long-range distances were incorporated in CYANA as upper and lower limit constraints to position the rigid bodies with respect to each other. Using CYANA, we calculated a raw ensemble of 20,000 models of which the 2,000 models with the lowest target function were selected (for details see Materials and Methods and Fig. S3). Subsequently, this ensemble was filtered for structural integrity and the individual conformers were refined using YASARA. Integrative ensemble fitting and contraction (i.e. reduction of the ensemble to a smaller number of conformers, while maintaining accurate representation of the experimental data) was performed with MMMx (52) to obtain a final ensemble of 25 conformers, each annotated with a distinct probability that provides a balanced fit to the 35 EPR distance distributions between protein sites and the three small-angle scattering curves (Fig. 2A). Altogether, this ensemble fits all the distance distributions, providing an overlap between experimental and back-calculated distance distributions between 0.396 and 0.845 with a geometric average of 0.662 (Fig. S5). A loss of merit of 0.332 upon integrating distance distribution and small-angle scattering restraints indicates that both sets of restraints are consistent with each other.

**Fig. 2.**
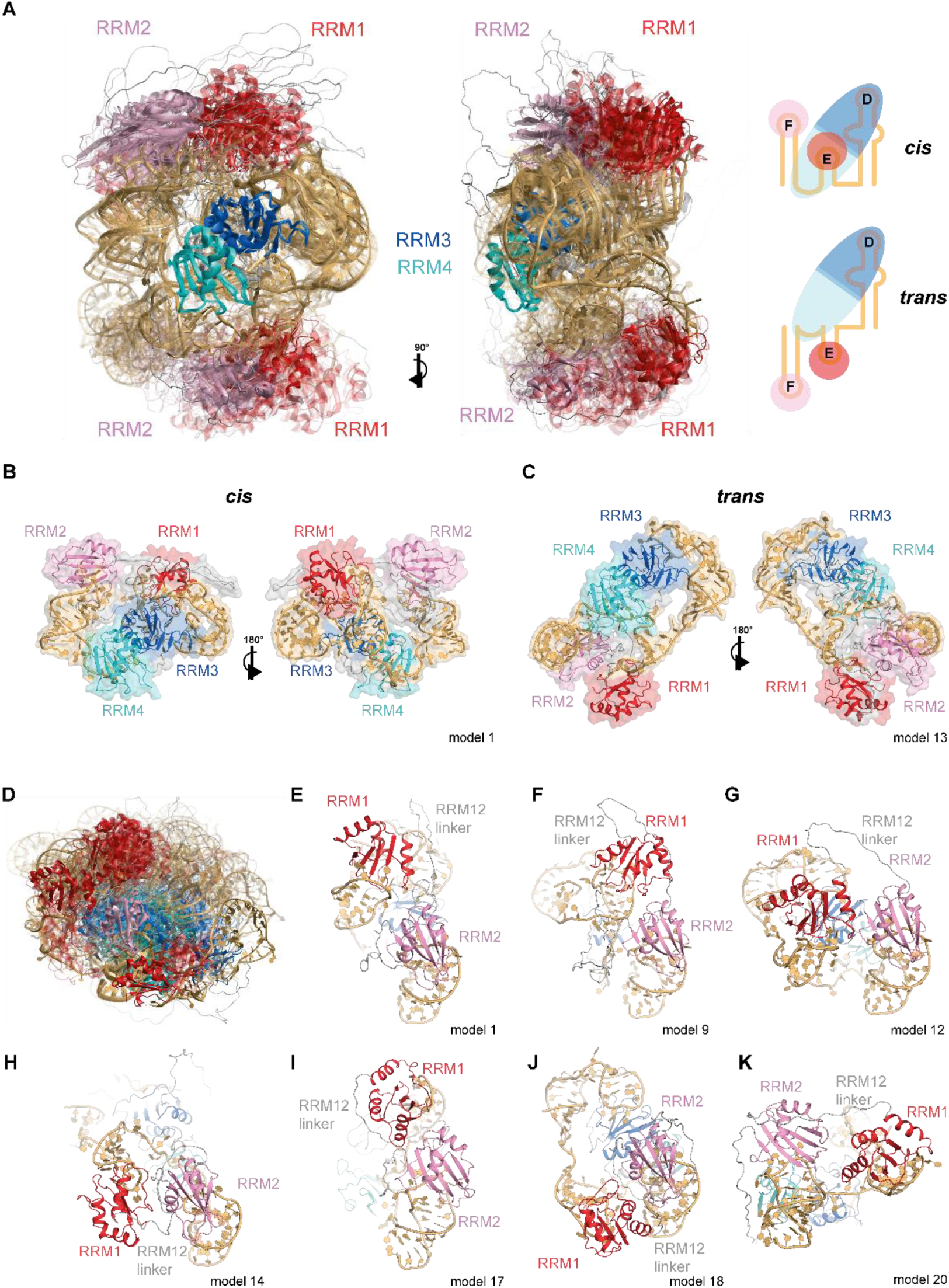
PTBP1 and EMCV-IRES-DtoF form a compacted complex with pronounced conformational flexibility. (A) Structural ensemble of the PTBP1/EMCV-IRES-DtoF complex in a conformer population-weighted visualization (opacity corresponds to the population of each conformer). The 25 models of the ensemble were superimposed on RRM34. Two views related by a vertical rotation of 90° are shown. SLE and SLF can be positioned in *cis* or *trans* with respect to SLD, illustrated in schemes on the right. (B) and (C) Conformers representative of the two subclasses (*cis* and *trans*) in two views. (D) Overlay of the ensemble on RRM2 illustrating the conformational flexibility of RRM1 with respect to RRM2. (E)-(K) Examples of conformers showing different ways RRM1 and RRM2 can be spatially close within the PTBP1/EMCV-IRES-DtoF complex.

### A compact structure spanning a large conformational space exposes the IRES RNA helices

The pairwise root mean square distance (rmsd) between the 25 conformers of the PTBP1/EMCV-IRES-DtoF ensemble varies from 2 to 30 Å over the entire complex (Fig. S6) reflecting the wide distance distributions measured for many spin pairs. Despite this apparent structural heterogeneity, the structural ensemble of the complex reveals very interesting features. The best way to illustrate the conformational space covered by this structural ensemble is to superimpose each conformer onto RRM34 (Fig. 2A). This representation reveals that RRM1 and RRM2 localize primarily on two sides with respect to RRM34. Based on the position of RRM1 and RRM2 relative to RRM34, the structural ensemble can be divided in two groups with a major form (13/25 conformers, with a 60% population by amounting the individual probabilities) and a minor form (12/25 conformers, with a 40% population). In the major form SLF and SLE are in *cis* with respect to SLD (Fig. 2B), while in the minor form, the two stem-loops are in *trans* with respect to SLD (Fig. 2C). In the *cis* conformation, RRM1 and RRM3 are close to each other, while in the *trans* conformation they are more distant. Importantly, despite being separated by a flexible linker, RRM1 and RRM2 are always within close proximity. Yet, RRM1 is not situated at a fixed position relative to RRM2 but occupies multiple sites with distinct orientations (Fig. 2D). Some of the determined RRM1 and RRM2 orientations can be explained energetically by the conformation of the inter-domain linker, molecular contacts involving the domains and the linker or the two domains and protein-RNA intermolecular interactions (see below) (Fig. 2E-K). Independent structural ensembles generated by either fitting only against DEER data (Fig. S7A) or after removal of the 25 conformers that were included in the best-fitted integrative ensemble (Fig. S7B), resulted overall in similar conformational ensembles composed of these two *cis/trans* families.

PTBP1 RRM34 interacts with EMCV-IRES-DtoF using the same interface and, remarkably, the same directionality, which we had previously identified when studying its binding to single-stranded RNA (53). Indeed, RRM3 binds the RNA at the 5’-end (SLD) and RRM4 further downstream (LinkEF) using the interaction surface near the small helix of the linker. This is seen in both the major and minor forms of the complex. However, only in the minor form, the *trans* arrangement brings the intervening RNA near the RRM34 inter-domain linker enabling additional protein-RNA interactions (Fig. 2C). As independent support for these contacts, we could identify protein-RNA cross-links between this region of the linker and the RNA (Fig. S8). The detection of these protein-RNA cross-links to U/UU fits with the proximity of the inter-domain linker with the region downstream of SLE (Fig. S8E). Strikingly, hydroxyl radical cleavage data (43) for the linker residue N432 support its position close to SLE (Fig. S9). The two main conformations originate from RRM4 binding to the linker between SLE and SLF, thereby restricting the mobility of the SLs, but still allowing SLD and SLE to be positioned in *cis* and *trans* as not all nucleotides of the LinkEF sequence are bound by RRM4. In the *trans* conformation, the RRM34 inter-domain linker seems to contribute to the stabilization of this form. Additional “weak contacts” between RRM1 and RRM2 then help further stabilizing most conformers of the complex (see below).

Importantly, despite the very strong compaction seen upon complex formation of the RNA and the protein (Fig. S7C), we still observe residual conformational flexibility (Fig. 2A). As indicated above, an unexpected and interesting feature of the PTBP1/EMCV-IRES-DtoF complex is the spatial proximity of RRM1 and RRM2, which is induced upon RNA binding. RRM1 and RRM2 do not strongly interact, resulting in RRM1 being in proximity to RRM2 but not at a fixed position (Fig. 2D). This proximity is facilitated by the 10 residues C-terminal to RRM1 that fold into an α3-helix upon binding to SLE, reducing the overall length and dynamics of the linker (38). Close examination of the 25 conformers revealed that RRM1 may favor conformations placing it close to RRM2 because the inter-domain linker is very hydrophobic and folds back as a hairpin, resulting in RRM1 facing the β-sheet of RRM2 (seen in models 1-7) (Fig. 2E). In other cases (models 11-12) (Fig. 2G), contacts between the α3-helix of RRM1 and the inter-domain linker between RRM2 and RRM3 brings RRM1 close to RRM2. In another subset (models 8-10), the α2-β4 loop of RRM1 interacts with the β3-α2 loop of RRM2 and the inter-domain linker makes contacts with the α1-α2 surface of RRM1, resulting in the β-sheets of both RRMs facing in the same direction (Fig. 2F). In a fourth subset (models 13-15), the RRM1-RRM2 proximity is mediated by the RNA, since SLE is sandwiched between both RRMs (Fig. 2H). In a fifth subset (models 16-17), α2 of RRM1 mediates contacts to the top part of RRM2 (Fig. 2I) and in one case (model 18), the two domains are arranged back-to-back (Fig. 2J). In a final subset (models 20-25), the two domains do not form any contacts and the inter-domain linker is elongated interacting with the RRMs (Fig. 2K). This illustrates the complexity of the conformational landscape of this type of protein-RNA complex, where weak interactions between folded domains, protein linkers and RNA are competing, resulting in a globally compact but still very dynamic structural ensemble.

Overall, a key aspect of this PTBP1-IRES structural ensemble is that protein binding results in a compacted RNP structure with reduced (compared to the free protein and RNA), but still pronounced conformational flexibility. Importantly, the four RRMs lie in the interior of the structure and expose the three conserved RNA stems to the outside (Fig. 2A-C). This conformation could explain the proposed chaperone role of PTBP1, since in exposing these RNA stems it presents the IRES structure for subsequent interactions with the ribosome or translation initiation factors as seen in IRES-ribosome structures (54-57) (see Discussion).

### RRM4-LinkEF interaction is crucial for a tight complex assembly and IRES activity

The RRM4-LinkEF interaction is central to the complex structure as it stabilizes the longest RNA linker (LinkEF) and thereby participates in the compaction of the complex. We therefore investigated the effect on the structural integrity of the complex by interfering with the RRM4-LinkEF interaction to test its functional importance (Fig. 3). Replacement of the pyrimidine tract in LinkEF with purines (Lmut in Fig. 3B) resulted in a complex with significantly broadened distance distribution between spin labels on RRM1 and RRM34 compared to the non-mutated complex (Fig. 3A), while the RNA secondary structure remained unchanged (Fig. S1F). Similarly, mutation of key amino acids for RNA-binding of RRM4 to alanine (H457A, R523A, K528A; referred to as RRM4ko) resulted in broader distance distributions between RRM4 and RRM1 or RRM2, respectively, compared to non-mutated PTBP1 (Fig. S10). These experiments confirm that RRM4 is crucial for the stabilization and compaction effect of PTBP1.

**Fig. 3.**
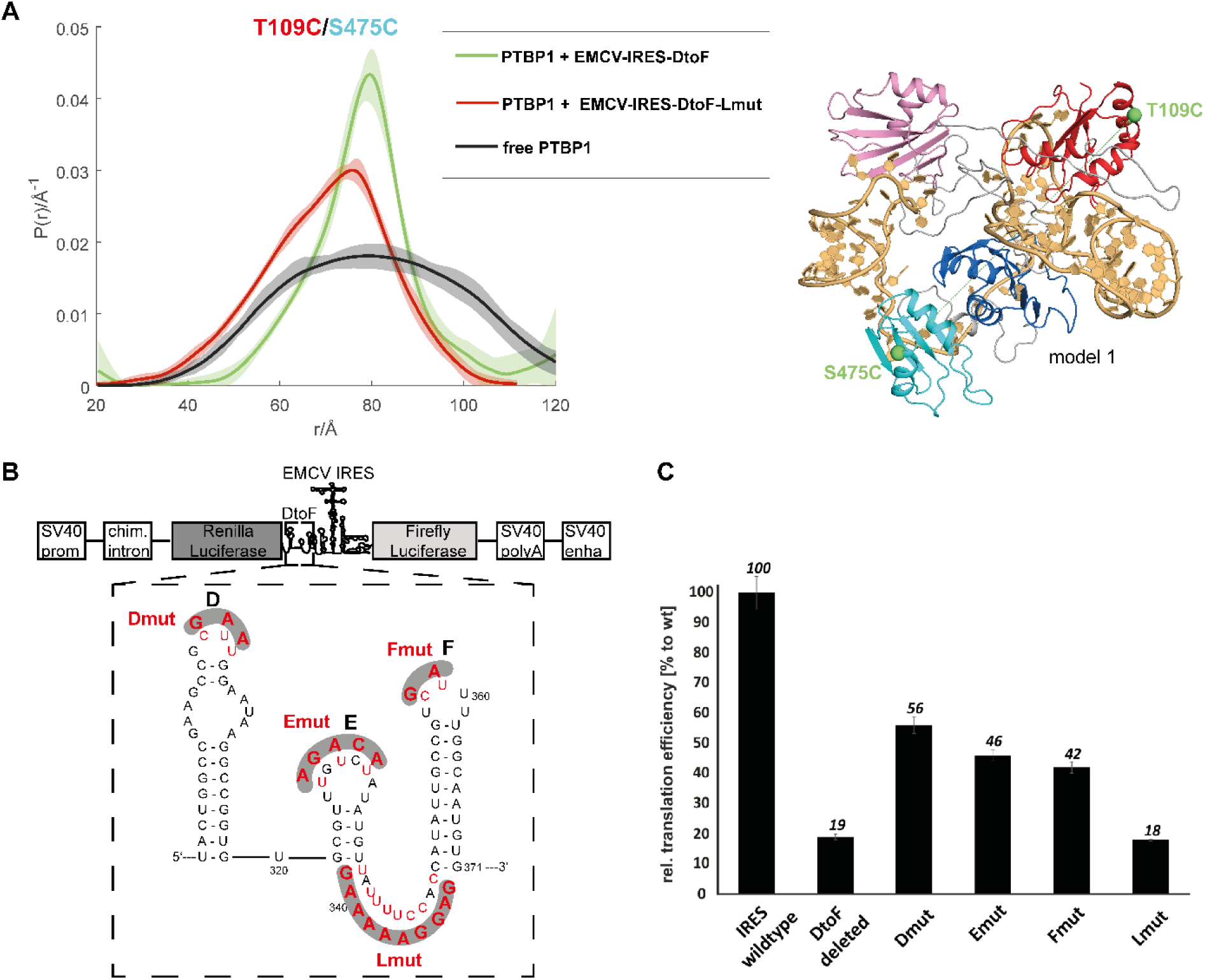
RRM4 binding is crucial for complex stabilization and translation enhancement. (A) Removal of the pyrimidine stretch in LinkEF (same mutation as shown in (B) for the Lmut Luciferase Reporter Assay construct) interferes with RNA binding of RRM4 and broadens the distance distribution measured between spin labels attached to RRM1 and RRM34 significantly. Compared to free PTBP1, the distance distribution narrows upon binding of EMCV-IRES-DtoF, but the mean distance does not shift substantially. This effect is less pronounced with the Lmut. (B) Constructs for the Luciferase Reporter Assay (top) with the deleted or mutated (pyrimidine to purine) sequence of EMCV-IRES-DtoF (bottom, only one mutation at a time). (C) Luciferase Reporter Assay given as IRES-mediated translation efficiency with respect to the WT-IRES. Remarkably, removal of RRM4-binding site (Lmut) results in the same activity loss as deletion of the whole EMCV-IRES-DtoF sequence, which links the importance of RRM4 binding for complex stabilization and rigidification with translational efficiency of the IRES.

To explore if altering the compaction of the complex would affect the biological functionality, we investigated the influence of the individual RRM-binding sites on IRES activity using a bicistronic luciferase reporter assay in HEK-293T cells (58). In this assay, Renilla Luciferase is translated 5’-cap-dependent and Firefly Luciferase is translated under the control of the EMCV-IRES, both from the same transcript. Thus, the ratio from Firefly Luciferase activity to Renilla Luciferase activity is a direct measure of IRES-activity (59). We first compared the activity of the EMCV-IRES wild-type (WT) (Fig. 3B,C) with a deletion mutant that lacks the entire EMCV-IRES-DtoF sequence (“DtoF deleted”). The deletion showed about a fifth of the WT activity (down to 19 ± 1.0 %), in line with previous reports that this part of the IRES is not essential but important for stimulating the IRES activity (33). Mutating the individual RRM binding sites on SLD, SLE or SLF by exchange of pyrimidines to purines led to a strong reduction down to 56 ± 2.7 %, 46 ± 1.8 % or 42 ± 1.8 %, respectively (Fig. 3B,C). Strikingly, mutating pyrimidines in the RRM4 binding site (LinkEF) to purines reduced the activity to 18 ± 0.4 %, similar to the deletion of the complete EMCV-IRES-DtoF. This strongly suggests that structural stabilization and compaction by RRM4 is crucial for EMCV-IRES activity.

Together these experiments revealed rather surprisingly that a flexible RNA linker, which connects two SLs, has an essential role in the biological activity of an RNA molecule. PTBP1 binds to the flexible RNA linker and remodels the overall RNA structure. The resulting compaction and the ordering of the RNA structure upon PTBP1 binding are crucial for the IRES-mediated translation initiation, potentially by facilitating the recruitment of the ribosome or interaction with translation initiation factors (28, 55–57). The overall conformation that exposes the RNA stems and the remaining conformational dynamics may allow the RNP to present a subset of conformations that can be recognized by the translation machinery through conformational capture. In contrast, without PTBP1 binding, the inherent flexibility due to lack of linker binding loosens the complex arrangement and therefore impedes translation initiation.

## Discussion

Here, we present unprecedented insights on the relationship between structure, dynamics and function of a multi-RBD protein-RNA complex by modelling its ensemble structure, which can lead to multimodal function. Since we took a new approach to determine the structure by integrating five structural methods and aiming at the best fit between the different structural constraints, two questions may arise: How accurate is the structural ensemble and what does it tell us about the biological function? Reassuringly, the presented structural ensemble is consistent with the hydroxyl radical cleavage that was performed by others (43) and thus this data can be seen as an independent support for the structural conformers (Fig. S9). Additionally, the identification of the *trans* conformation helped us explain, retrospectively, the cross-links we identified earlier (39) between the linker of RRM34 and the RNA (Fig. S8). Functionally, our results rationalize the proposed roles of PTBP1 as RNA chaperone and ITAF in the context of IRES-mediated translation initiation. In recent years, a few structures of IRES elements bound to ribosomes have been determined (54–57, 60), however, in all these cases, the IRES RNA is stabilized exclusively by pseudoknots and RNA tertiary interactions, although most viral and cellular IRESs require protein ITAFs for optimal translation activity. Our structural work provides evidence that PTBP1 strongly compacts the RNA and stabilizes the RNA fold into discrete conformations. As a result of PTBP1 binding, the RNA helices of SLD, E and F are exposed to the solvent for further interactions with the translation machinery, consistent with what was found in protein-free IRESs bound to ribosomes (54–57, 60). Interestingly, in particular the sequence of the solvent-exposed SLD containing a purine-rich internal loop, is conserved suggesting that the structure and the sequence of the exposed IRES element may be recognized by the ribosome or additional factors (Fig. S1). In the context of EMCV-IRES, the SL J-K are bound by the HEAT-1 domain of eukaryotic initiation factor 4G (eIF4G), which require a pre-organization to position protein-recognition (61). Interestingly, a second PTBP1 molecule has been shown to bind to this region of the EMCV-IRES and thereby may contribute to this organization, which has been shown to be common in type 2 picornavirus IRESs (43, 62). The compaction and stabilization effect was not necessarily expected because PTBP1 itself is a very dynamic protein containing four folded domains separated by flexibles linkers and previous studies on the shape of the free protein have established the view that PTBP1 is present in solution as an elongated particle (36, 48). A number of features unique to PTBP1 make the protein able to fold EMCV-IRES-DtoF into a structured RNA: First, the four RRMs separated by flexible linkers allow binding to separate RNA sequences within the IRES. The observed RNA structure stabilization through PTBP1 binding explains also previous data showing that mutations of individual RRMs do not disrupt RNA binding, but still result in strong reduction of translation efficiency (43). Second, the tight interaction between RRM3 and RRM4 brings distant RNA sequences into close proximity. Our structural and functional study identifies the interaction of RRM4/LinkEF as a major element for structuring the core of PTBP1/EMCV-IRES-DtoF. An increase in flexibility of the complex upon mutation of the RRM4 binding site in LinkEF coincides with a dramatic reduction in translation efficiency. Third, binding of RRM1 to SLE results in a shortening of the linker between RRM1 and RRM2 (38). Fourth, there is a close interaction between RRM1 and RRM2 that is mediated by several sets of interactions, and not a unique one. This last feature is primarily responsible for the residual conformational flexibility seen in the final structural ensemble. Conformational flexibility was also seen in protein-free IRES structures (54–57, 60), so it seems to be a common structural feature of IRESs. This flexibility may help with the initial recognition by the ribosome and also for the IRES to adapt its structure during the different phases of translation that are accompanied by structural dynamics of the ribosome. In agreement with this, recent *in vivo* single-molecule experiments showed that EMCV-IRES transitions between translationally active and inactive RNA states (63). Such RNA remodeling capability of PTBP1 is also likely to be important for splicing regulation, consistent with reports supporting the role of RRM4, and the interaction with RRM3 for splicing activity (53, 64–66). Thus, compaction and stabilization of the RNP complex is not restricted to IRES-mediated translation initiation, but a general mode of action of PTBP1.

The structural details presented here also provide general insights into how RBPs interact with RNAs. To our knowledge, the only other structure of a multi-RRM containing protein with more than two RRMs in complex with a natural RNA is the crystal structure of the spliceosome-specific RNA chaperone Prp24 bound to U6 snRNA (67). Both Prp24 and the U6 snRNA experience large structural rearrangements upon complex formation, and the crystal structure has a rigid interlocked topology. In this case, only three of the four RRMs interact with RNA (RRM2-4) and RRM2 interacts extensively with RRM1 and RRM4, while the inter-domain linker between RRM3 and RRM4 folds into two α-helices upon RNA binding. Therefore, PTBP1 and Prp24 use completely different mode of actions although both act as RNA chaperones. Importantly, while Prp24 has a single RNA binding substrate, PTBP1 has thousands of RNA binding targets being involved in almost all post-transcriptional gene regulatory processes (68). Therefore, it is maybe not surprising that complex formation of PTBP1 results in a discrete number of complex structures and not a single one. Moreover, it is likely that it would be energetically more favorable for a RNP complex to have a higher degree of flexibility and thereby maintaining a higher conformational entropy compared to a rigid structure, especially considering the large entropic cost associated with the binding between two very flexible molecules. The preservation of conformational dynamics in the bound state may also facilitate the recruitment of additional binding partners and allow them to exert additional influence on the RNAs conformational landscape, required to attain functional conformations.

We expect that the approach presented here will prove valuable for future structural characterizations of RNP complexes involving the numerous RBPs in the human genome that, similarly to PTBP1, are composed of multiple RBDs separated by flexible linkers. RNA binding may result in several, substantially different conformations and could be vital to their function. Many RNP machineries rely on major structural rearrangements as seen in the different steps catalyzed by the spliceosome or the ribosome (19, 20). Our approach that combines short and long-range distance constraints from NMR, MS and SAS, as well as, distance distributions from EPR, allows to determine a structural ensemble covering the full conformational space covered by such dynamic RNPs. Distance distributions from EPR experiments inform on the width of conformational distributions and thus represent a basis for integrative structure modeling of large systems, provided that three-dimensional models of the structurally well-defined building blocks already exist. The recent advances in AI-driven structure predictions will aid the availability of such building blocks (69). Integrating NMR and EPR data, protein-RNA cross-linking restraints and SAS curves holds much promise as a strategy to determine ensembles of rigid-body arrangements and even protein-RNA condensates, to provide new insights into the relationship between structure, dynamics and function of RBPs.

## Materials and Methods

### Site-directed mutagenesis

Positions for site-directed spin labeling (SDSL) on the protein were chosen based on simulating the spin label attachment to existing NMR solution structures of individual RRMs of PTBP1 (8) using the software package Multiscale Modelling of Macromolecular systems (MMM) (70). For inter-domain EPR distance measurements, we chose amino acids in the RRMs that are predominantly located within the α-helices, with geometrical similarities and related chemical properties compared to cysteine. Positions in the peptide linker regions were selected closed to the C-terminus of RRM1 and RRM2, close to the N-terminus of RRM3 or in the center of the linker.

### Protein expression and purification

Protein expression and purification of PTBP1 and individual RRMs was performed as described previously (39, 47). For distance measurements > 5.5 -6 nm, expression was performed in 95-99 % D_2_O using minimal M9 medium followed by same purification protocol. Samples for NMR and SANS experiments were finally transferred to 10 mM NaPO_4_, pH 6.5, 20 mM NaCl buffer with 1-2 mM DTT by size exclusion chromatography and dialysis.

### RNA preparation

*In vitro* transcription of EMCV-IRES-DtoF and of individual SLs was performed as described previously (39, 47). For spin labeling of RNA, respective RNA segments were removed by RNase H cleavage and chemically synthesized (Dharmacon) thio-uridinated RNA segments were ligated after spin labelling similarly as described before (47, 49, 71).

### Site-directed spin labeling of protein and RNA

Spin labeling protocols used for all PTBP1 mutants have been described elsewhere (47). For SDSL of PTBP1, we used predominantly MTSSL ((1-oxyl-2,2,5,5-tetramethylpyrroline-3-methyl) methanethiosulfonate, Toronto Research Chemicals) or IAP (3-(2-iodoacetamido)-proxyl, Sigma Aldrich). For RNA SDSL, different uridine positions in the loop or linker regions in EMCV-IRES-DtoF were selected and the respective short oligonucleotides containing 4-thiouridine were spin labeled and ligated to the full-length RNA following earlier published protocols (47, 49).

### PTBP1/EMCV-IRES-DtoF complex formation

Complex formation and purification was performed essentially as described before (39). In brief, EMCV-IRES-DtoF was diluted in all cases to approximately 4 µM in 5 mL low salt buffer (10 mM NaPO_4_, 20 mM NaCl, pH 6.5). Concentrated doubly-labeled PTBP1 was mixed with the RNA in a molar ratio of 1:1.2 and the complex was then purified by size-exclusion chromatography using a Superdex 75 column (Cytiva). Complex samples were then buffer-exchanged into D_2_O and concentrated to approximately 50-100 µM. For DEER distance measurements, complex samples were diluted 1:1 (v/v) with d_8_-glycerol (Sigma Aldrich) and 30 µL sample solution was filled into 3 mm quartz tubes. Complex formation was ensured by native RNA polyacrylamide gels.

### Bicistronic reporter assays

Bicistronic pRemcvF plasmids were a generous gift from Prof. Dr. A. Willis (MRC Toxicology Unit, Leicester, UK) (72). EMCV-IRES mutants were generated using standard site directed mutagenesis procedures. HEK-293T cells were maintained in Dulbecco’s modified Eagle’s medium supplemented with 10% heat-inactivated fetal calf serum (GibcoBRL) and antibiotics. For IRES activity assays, a 24 well plate was seeded with 500 µL cell suspension (approx. 350,000 cells/mL). After 24 hours growth, the medium was exchanged to medium without antibiotics. Then, cells were transfected with 500 ng plasmid/well and a final volume of Lipofectamin 2000 (Thermo Fisher) of 1 µL/well in a total volume of 100 µL OPTI-MEM (Gibco) medium/well. Cells were incubated for 24 hours and lysed using the Dual-Luciferase Assay lysis buffer supplemented with Roche Complete Protease Inhibitor (Roche, incubation of 10 min on a shaker). Lysates were cleared by centrifugation (13,500 rpm, 4°C). 5 µL of lysate/well were pipetted into a 96-well ELISA plate, 25 µL of Firefly-Luciferase substrate solution were added and the measured after 15 seconds shaking in the plate reader. Then, 25 µL of Stop-N-Go solution were added and samples were measured to access the Renilla-Luciferase activity. All IRES assays were performed in technical triplicates of biological triplicates and measured on a Berthold MicrolumatPlus luminometer. IRES activity is determined by normalizing the Firefly-Luciferase activity to the Renilla-Luciferase activity whereas the activity of the WT-plasmid is set to 100%. Error bars represent the errors as estimated by error propagation.

### CLIR-MS/MS

Mass spectrometry data from previous CLIR-MS/MS analysis of the PTBP1-IRES complex (39) was reanalyzed with an optimized parameter set containing more neutral loss products, to increase the number of identified protein-RNA cross-links. Mass spectrometry data files (Thermo Fisher RAW format) corresponding to the uniformly labeled EMCV-IRES RNA experiment were retrieved from ProteomeXchange via the PRIDE partner repository (dataset identifier: PXD005566) and converted to mzXML using msconvert.exe (ProteoWizard msConvert v.3.0.9393c) (73). The files were searched using xQuest (v2.1.5, available for download from https://gitlab.ethz.ch/leitner_lab/xquest_xprophet) (74, 75) against a database containing only the target PTBP1 protein sequence, with expected light-heavy RNA adducts defined as monolinks. Adducts with lengths of 1-4 nucleotides were considered. In addition to the whole RNA adduct, the following neutral losses were considered: -H2O, -H4O, -HPO3, -H3PO3, -H2, -HPO3+H2O, -HPO2, -H4O2, -H5PO5. Further details about the expanded parameter set can be found elsewhere (76). All amino acids were set as cross-linkable. Delta masses for each adduct were defined depending on the nucleotide sequence composition, as described previously (39). Further parameters used for xQuest searching: delta mass tolerance: +/- 15 ppm, retention time tolerance: 60 s, enzyme = Trypsin, maximum missed cleavages = 2, MS1 mass tolerance = 10 ppm, MS2 mass tolerance = 0.2 Da. Identifications with an ld.Score > 20 (according to the scoring scheme described previously (74)) were considered. Further processing and visualisation was undertaken using custom Python 3.7.1 scripts. The reanalysed mass spectrometry proteomics data have been deposited to the ProteomeXchange Consortium (http://proteomecentral.proteomexchange.org) via the PRIDE partner repository (77) with the dataset identifier PXD034894.

### NMR spectroscopy

All NMR-experiments were recorded on Bruker Avance III 500, 600, 700 or 900 MHz spectrometers equipped with cryo-probes and on a Bruker Avance III 750 MHz spectrometer with a room temperature probe. NMR protein spectra (free proteins and protein-RNA complexes) were acquired at 313 K and 323 K, imino ^1^H^1^H-NOESY spectra were recorded at 278 or 283 K. Imino-^1^H^15^N-TROSY spectra were recorded at 278 K and 293 K. We processed spectra with Topspin 2.1 or Topspin 3.0 and analyzed in Sparky 3.0 (78). ^1^H-^1^H imino assignments of the RNA were achieved by standard methods (79, 80). Imino resonances of SLE were assigned using conventional jump-return NOESY facilitated with SMT experiments at 278 and 275 K (81, 82). Non-exchangeable protons were assigned using sequential walk in NOESY spectra recorded at 298 K and in 100% D_2_O. Assignment of other hydrogens and heteronuclei were obtained using intra-nucleotide assignment experiments including, aromatic HSQC, CT HSQC of sugar resonances, TROSY HCCH COSY for adenine H2-H8 assignment, HCCH COSY for H5-H6 correlations of pyrimidines, and finally HCN and HCNCH experiments. Assignments of protein resonances were obtained using conventional triple resonance experiments, HCCH-TOCSY and 3D NOESY-HSQCs. Intermolecular NOEs were identified in ^13^C,F3-filtered HSQC-NOESY experiments.

### EPR spectroscopy

Labeling efficiencies and spin label tumbling regimes of spin labeled protein and RNA samples were determined by CW EPR spectroscopy at ambient temperature. Experiments were performed at X band (9.5 GHz) using a Bruker Elexsys E500 spectrometer with a Bruker super-high Q resonator ER4122SHQ and spectra were recorded with a field modulation of 100 kHz, a modulation amplitude of 1 G, a time constant of 10.24 ms, a conversion time of 40.96 ms and an attenuation of 25 dB of 200 mW incident microwave power. Samples were filled into glass capillaries (BLAUBRAND®) with a diameter of 1.5/0.9 mm (outer/inner). Labeling efficiencies were determined by digital double integration of the EPR spectra and comparing them to free nitroxide radical IAP of standard concentration. Changes in the spin label mobility could be detected by amplitude reduction of the low- and high-field component of the nitroxide spectra which indicated successful spin labeling.

To obtain EPR distance restraints, four-pulse DEER experiments (46) were performed at a home-built high-power Q-band spectrometer (35 GHz) with 200 W microwave power, a Bruker ElexSys acquisition system (E580) and a home-built TE001 pulse probe (83). Temperature stabilization during all measurements was ensured by a He-flow cryostat (ER 4118CF, Oxford Instruments) and a temperature control system (ITC 503, Oxford Instruments). All free protein and complex measurements were carried out at 50 K because this temperature corresponds to the optimal conditions for nitroxide radicals with respect to their longitudinal and transverse relaxation. Pump pulses were always set to 12 ns and detection pulses to 16 ns. The first inter-pulse delay τ_1_ was set to 400 ns, τ_2_ was set according to the expected distance and its required DEER trace length for suitable background correction. Time increment *t* was also adapted to the length of the τ_2_ as described earlier (47). The pump pulse was always applied on the maximum field position of the nitroxide spectrum, whereas detection pulses were applied by an offset of approximately 100 MHz. Samples which were diluted 1:1 (v/v) with d_8_-glycerol (Sigma Aldrich) were shock frozen with liquid nitrogen and measured in this state.

### Analysis of DEER data

For processing all DEER data we used the software package DeerAnalysis (84), version 2022 in automated mode. When an artefact arose from overlap between the excitation bands of pump and detection pulses (84, 85), up to 15% of the data was cut at the end. For background correction we used a dimensionality of 3, corresponding to a homogenoeus three-dimensional distribution of complexes in the sample as expected for soluble proteins. Automated comparative analysis computes distance distributions by neural network analysis with DeerNet (86) and Tikhonov regularization (87) with DeerLab in a single step with background correction (88) and provides 95% confidence intervals for both distributions. Unless otherwise indicated, we report the mean of the two distributions and confidence intervals that include uncertainty due to model bias. The mean distances (〈r〉) and standard deviations (σ(r)) of the distance distributions were implemented as upper and lower limits in CYANA modeling (see below).

### Small angle scattering experiments

Small angle neutron scattering experiments were recorded at the SANS-I and SANS-II facility, Swiss Spallation Neutron Source, SINQ, Paul Scherrer Institute, Switzerland. Scattering densitiy matching points for RNA and protein with respect to the D_2_O/H_2_O ratio of the buffer (10 mM NaPO_4_, pH 6.5, 20 mM NaCl, 2 mM DTT) were determined by contrast variation and extrapolation of I_0_ to be 66% D_2_O for RNA, and 42% or 112% for protonated or deuterated protein, respectively. Free protein and RNA were recorded at a wavelength of the neutron beam of 6 Å, a collimation of 6 m and a detector distance of 2 m and 6 m.

For both, SANS and SAXS experiments, complexes were reconstituted as described previously (39) and dialyzed for 24 h against suitable buffers. Each reference cuvette was filled with the corresponding dialysis buffer. Complexes were measured at SANS-I at a wavelength of 4.5 Å and either a collimation of 11 m and a detector distance of 11 m or a collimation of 3 m and a detector distance of 2 m. Scattered neutrons were detected using a two-dimensional 96×96 cm^2^ detector with a pixel size of 0.75 cm. The cross section of the collimator was 50×50 cm^2^ and the effective sample diameter was 0.8×1.5 cm^2^. At SANS-II, the selected wavelength was 4.9 Å, and collimation and detector distance were 2 m and 1.2 m or 4 m and 4 m, respectively. The detector had a diameter of 64 cm with 128×128 pixels. Reduction and analysis of SANS data was performed with the program BerSans (89) and visualized using Primus QT (90) from the ATSAS program (91). Beamline specific correction factors for data reduction are 1.338 and 1.284 for data recorded at a wavelength of 4.5 and 6 Å, respectively. SAXS experiments were performed on the liquid samples using a Rigaku MicroMax-002 microfocused beam (4 kW, 45 kV, 0.88 mA). The Cu Kα radiation (λ_CuKα_ = 1.5418 Å) was collimated by three pinholes (0.4, 0.3, and 0.8 mm) collimators. The scattered X-ray intensity was detected by a two-dimensional Triton-200 X-ray detector (20 cm diameter, 200 mm resolution) for SAXS. SAXS detectors cover an effective scattering vector range of 0.1 nm^-1^< q < 4 nm^-1^, where q is the scattering wave vector defined as q = 4π sin θ/λ_CuKα_, with a scattering angle of 2θ. The SAXS samples were measured in 2 mm quartz X-ray capillaries (purchased from Hilgenberg).

### Modeling with CYANA

For the integrative structure modeling of PTBP1/EMCV-IRES-DtoF (Fig. S3) we used the previously determined CLIR-MS/MS derived models of PTBP1 RRM1, RRM3 and RRM4, as well as, the NMR-derived solution structure of RRM2 in complex with the respective RNA sites as rigid bodies (39). However, as CYANA does not allow to load protein-RNA complexes as input structures, protein and RNA were loaded separately. Based on the imino-^1^H^1^H-NOESY signals and structure prediction by McFold/McSym, we modeled RNA-stems using standard A-form RNA-helix angles and Watson-Crick base pair restraints^45^. We merged the protein-models (only first state of the bundle) and RNA models in one PDB file (numbering was adjusted to fit all models on a single chain as required for CYANA) and used the regularize macro of CYANA to generate a similar (but not identical) bundle of PTBP1 input structures (RMSD < 0.3 Å). Subsequently, the definition of rigid bodies in CYANA for protein domains was achieved using the command “angle fix”. To preserve the RNA secondary structure, Watson-Crick base pair restraints were enforced in the stem, while RNA loops and bulges were kept flexible. These regularized rigid bodies were used as input for a CYANA calculation. The linkers between these rigid domains were generated with random starting conformations by CYANA. In these calculations, intermolecular restraints that were adapted from previous work (39), were used to define protein-RNA contacts as observed in the individual sub-complexes.

To include EPR distance distributions in CYANA, we generated RRM models with 50 rotamers of the corresponding spin labeled cysteine and calculated the geometric center of the radical positions. This average position was represented by a dummy glycine-Cα that was fixed by three or four strong restraints to the domain or linker-peptide, respectively (upper limits and lower limits in CYANA, ±0.05 Å of geometric center). A final set of 35 EPR distance restraints were implemented as upper and lower limits corresponding to the mean plus or minus one standard deviation of the distance distribution, respectively.

As CYANA does not include electrostatics, we used artificial lower limits of 6 Å between RNA-phosphate groups with corresponding upper limits set to 200 Å. The latter distance is larger than D_max_ determined by SAS experiments and, thus, does not artificially compact the RNA.

Using these input rigid bodies (protein domains and RNA fragments) in combination with the short and long range intermolecular restraints, we executed a simulated annealing protocol with 30,000 steps and calculated 20,000 models (20 x 1,000 with different seeds). Out of these 20,000 models, the 10% energy best structures (lowest target function) were selected for further refinement.

### Structure refinement

All pseudoatoms from the CYANA output were removed, it was split into two chains (protein and RNA) and the RNA was renumbered by a customized script. Afterwards, models with flexible peptide linkers that are threaded through RNA or contain C4’-O4’ bond lengths in RNA sugars longer than 1.6 Å were discarded. These tests discard between 2 and 8 models per run (out of 100 models). The models, which passed both tests, were refined by YASARA using a standard script. The script solvates the model with water, runs restrained MD, and optimizes the structure that way. The resulting structure remains close to the input structure, while removing clashes, and fixing wrong bond lengths, wrong bond angles, and wrong torsion angles.

The individual output models of YASARA have different protonation states of RNA, however, models in a PDB file must have the same number of atoms. Therefore, all protons that did not exist in all structures of the first run were removed to obtain consistent models with minimal protonation. Models with fewer protons than the first model, which usually indicates problems in RNA geometry, were discarded. Once again models were checked for flexible peptide linkers threaded through RNA. This is necessary because some CYANA models were very strained and their structures changed so strongly during refinement that such threading occurred even if it did not exist in the input model of the YASARA refinement. In the case of threading after YASARA refinement, models were discarded. Similarly, sugar bonds were checked again. When trying to remove strain, YASARA sometimes breaks a C4’-O4’ bond, because this is less costly in terms of energy than the alternatives. All models with altered sugar bonds were discarded. The set of all YASARA-refined CYANA models that passed these tests contained 766 conformers, and furnished the raw ensemble for ensemble refinement.

In total, 35 EPR (distance distribution) restraints, the SAXS curve and two SANS curves (66% D2O, 1.2 and 4 m detector distance) were used for refinement. An additional ensemble was refined only with the EPR restraints.

Fitting was done in blocks of 100 conformers from the raw ensemble of 766 conformers with an algorithm searching for the global minimum by adjusting their populations (52). In each such fit, all conformers with less than 1% of the population of the most populated conformer were discarded. Many populations were driven to exactly zero. Thus, after fitting one block of 100 conformers, a preliminary ensemble of much less than 100 conformers, together with corresponding populations, was obtained.

This preliminary ensemble was topped up to obtain a new block of 100 conformers, by adding conformers from the raw ensemble that were not part of the first block of 100 conformers. The new block was fitted to obtain another preliminary ensemble with less than 100 conformers. This step was repeated multiple times, using up all conformers of the raw ensemble, with the last block usually containing less than 100 models.

In each block, the following fitting steps were performed. First, the block was fit only to the EPR restraints. This provides the best overlap deficiency (52) with the conformers of this block by assigning populations (e.g. OD1). Overlap for a single distance distribution restraint corresponds to the common area below the experimental and simulated distance distributions, whereas the total area of each distribution is normalized to 1. It ranges between 0 for completely distinct and 1 for identical distributions. Overlap deficiency for the set of all distance distribution restraints ranges between 0 and 1, with 1 meaning no overlap for at least one restraint and 0 meaning perfect overlap for all restraints. Second, the block was fit to only the three small-angle scattering (SAS) curves by minimizing the sum of the χ^2^ values for all SAS curves. This provides the best sum of χ^2^ values that can be achieved for the conformers in this block by assigning populations (e.g. SCS1). Finally, the block was fit to EPR and SAS restraints simultaneously. In this fitting step conformers of the block are selected, and populations are determined. Fit quality of EPR and SAS restraints are balanced. For each trial set of populations, the mean overlap deficiency (OD2) and the sum of χ^2^ values for the SAS curves (SCS2) are computed. Because this cannot be better than OD1 in this block, OD2 ≥ OD1. Likewise, it cannot be better than SCS1, i.e. SCS2 ≥ SCS1. Hence, (OD2/OD1 + SCS2/SCS1)/2 ≥ 1. The minimum of 1 can only occur if the same set of populations provides the best fit for EPR restraints as well as the best fit for SAS restraints. If the restraints are inconsistent or the set of input conformers is not good enough, the value will be substantially larger than 1. In order to get a transparent measure, we defined the loss of merit, (OD2/OD1 + SCS2/SCS1)/2 - 1. The loss of merit is zero, if both restraint sets are best fitted by the same set of populations. If it is one, the two figures of merit (overlap deficiency and sum of χ^2^ values) have approximately doubled. If this happens, the subsets of restraints are inconsistent or the set of conformers is very poor. Values for the loss of merit of 0.2 to 0.4 are quite normal (92). The loss of merit for PTBP1/EMCV-IRES-DtoF ensemble refined with both DEER and SAS data is 0.332 and thus indicates overall consistent restraint sets and supports the integrity of the CYANA raw ensemble.

## Acknowledgments

This work was supported by the SNF project 31003A-149921 and 31003A-170130 and the SNF-NCCR RNA and Disease (to F.H.-T.A.). C. G., M. Y. and G. J. thank ETH Zürich for the financial support (Grant No. 0-2066-14). G. J. and F.H.-T.A. also thank the Sinergia grant CRSII5-170976 for financial support. G.D. was supported by a PhD fellowship from the Boehringer Ingelheim Fond. T.d.V. and C.P.S. were funded by an ETH Research Grant (grant ETH-24 16 2). T.J.W would like to thank the ThinkSwiss Scholarship for funding.

## Author contributions

G.D., C.G., M.Y., G.J and F.H.-T.A. designed the project. G.D, C.G, T.d.V., C.M. and T.J.W. prepared protein and RNA samples. G.D., M.N. and F.F.D. performed NMR data collection and analysis. C.G., M.Y., T.J.W. and G.J. performed EPR data collection and analysis. G.D., E.D, S.B., R.M. and J.K. performed SANS/SAXS data collection and analysis. T.d.V., C.P.S., R.A and A.L. performed re-analysis of CLIR-MS/MS data. G.D., T.d.V. F.H.-T.A. and G.J. performed integrative structural modeling and analysis. E.F. performed bioinformatics work. T.d.V., F.H.-T.A., G.D., C.G. and G.J. wrote the manuscript with contributions and critical feedback from all the authors.

---

**Fig. S1.**
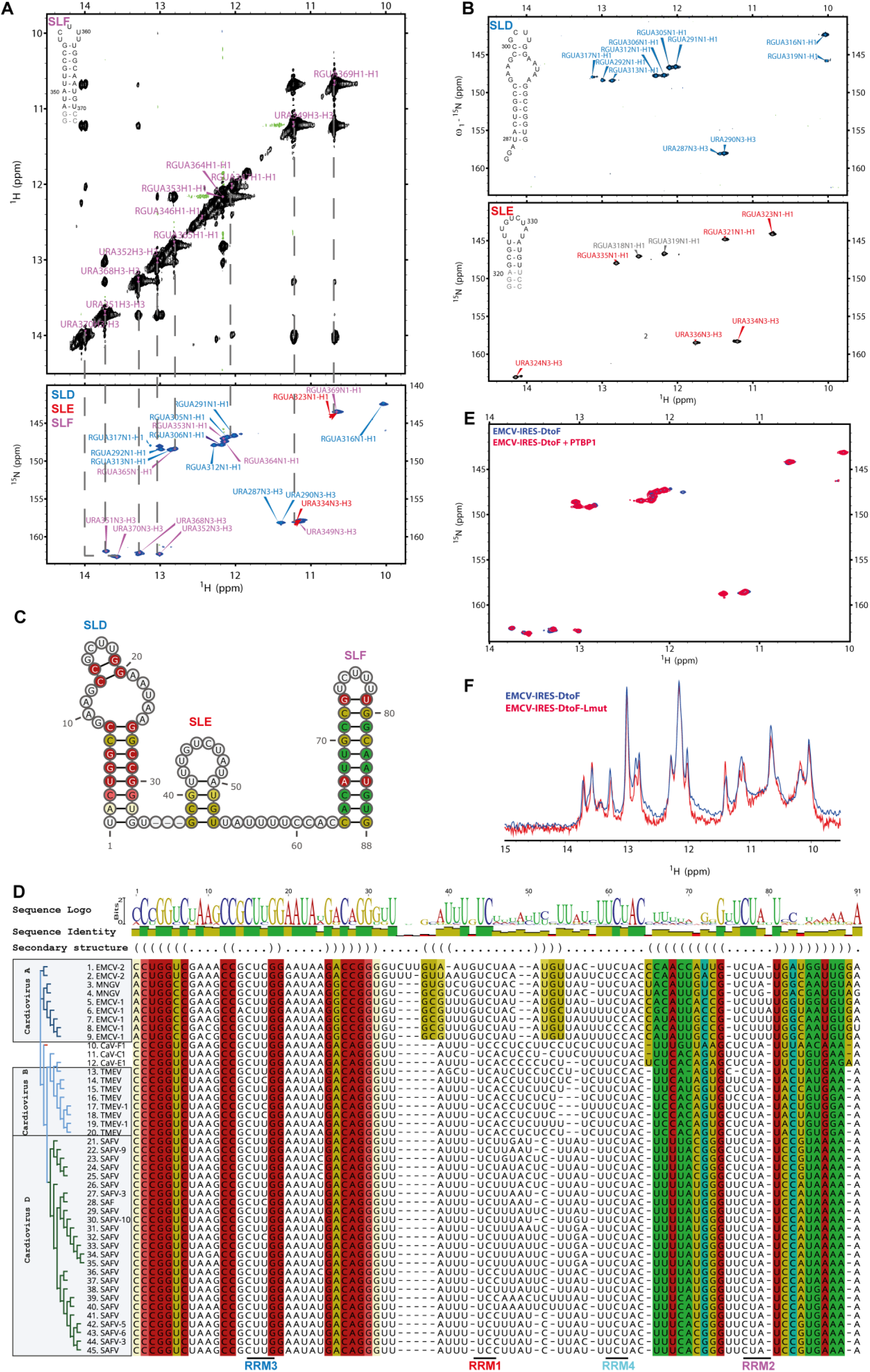
Secondary structure probing and phylogeny of EMCV-IRES-DtoF. (A) Assignment of imino proton resonances using a 2D ^1^H-^1^H NOESY experiment (top) and standard “sequential walk” exemplified on SLD. Signals could be readily transferred to the 2D ^1^H-^15^N HSQC of the full-length RNA (bottom). (B) Transfer of imino proton assignments to the corresponding 2D ^1^H-^15^N HSQC of each SL served as validation and comparison with the typical imino resonance shifts of A-U, G-C and G-U base-pairs and enabled unambiguous assignment. The match of the imino proton resonances recorded on the individual stem loops as well as on the complete RNA sequence showed that the proposed secondary structures of the RNA are present in solution. (C) Consensus secondary structure for these RNA elements in the genome of cardioviruses A (EMCV-1, EMCV2 and MNGV). The base-pairs are color-coded on the predicted minimum free energy structure according to base-pair probabilities from the multiple sequence alignment (MSA), warm colours mean high probability and cold colours mean low probability. (D) MSA from unique sequences in the 5’ UTR of all cardioviruses. Sequence logo, sequence identity (conservation) and secondary structure in dot-bracket annotation are shown on top. Columns of aligned RNA bases are colour-coded to correspond with the consensus RNA secondary structure. Complete genome sequences were retrieved from NCBI database using the key word “Cardiovirus”. An MSA was built using MAFFT software (93) and it was trimmed to include the SLD, E and F in the 5’ UTR of the cardioviruses genome only. RNAalifold software (94) was used to obtain the consensus secondary structure and its predicted free energy. Sequence logo and sequence identity were computed using geneious 2022.1.1 software. Sequences used for the alignment are listed in Table S1. The binding sites of the RRMs of PTBP1 according to (39) are indicated below the alignment. (E) 2D-^1^H-^15^N-TROSY spectra of free EMCV-IRES-DtoF and bound to PTBP1. Imino ^1^H-^15^N signals of the free EMCV-RNA (blue) and the PTBP1-EMCV-complex (red) remain at the same chemical shift in proton and nitrogen. This strongly indicates that the local RNA stem-loop structures remain unchanged upon protein binding. The NMR spectra were recorded at 288 K. (F) Overlay of 1D-^1^H-spectra recorded on EMCV-RNA with pyrimidine-to-purine mutations in LinkEF (Lmut, red) and wild-type (WT, blue) RNA. The signature of the imino-protons remains the same thus the secondary structure elements of the RNA were not affected by the mutations of the single stranded LinkEF region.

**Fig. S2.**
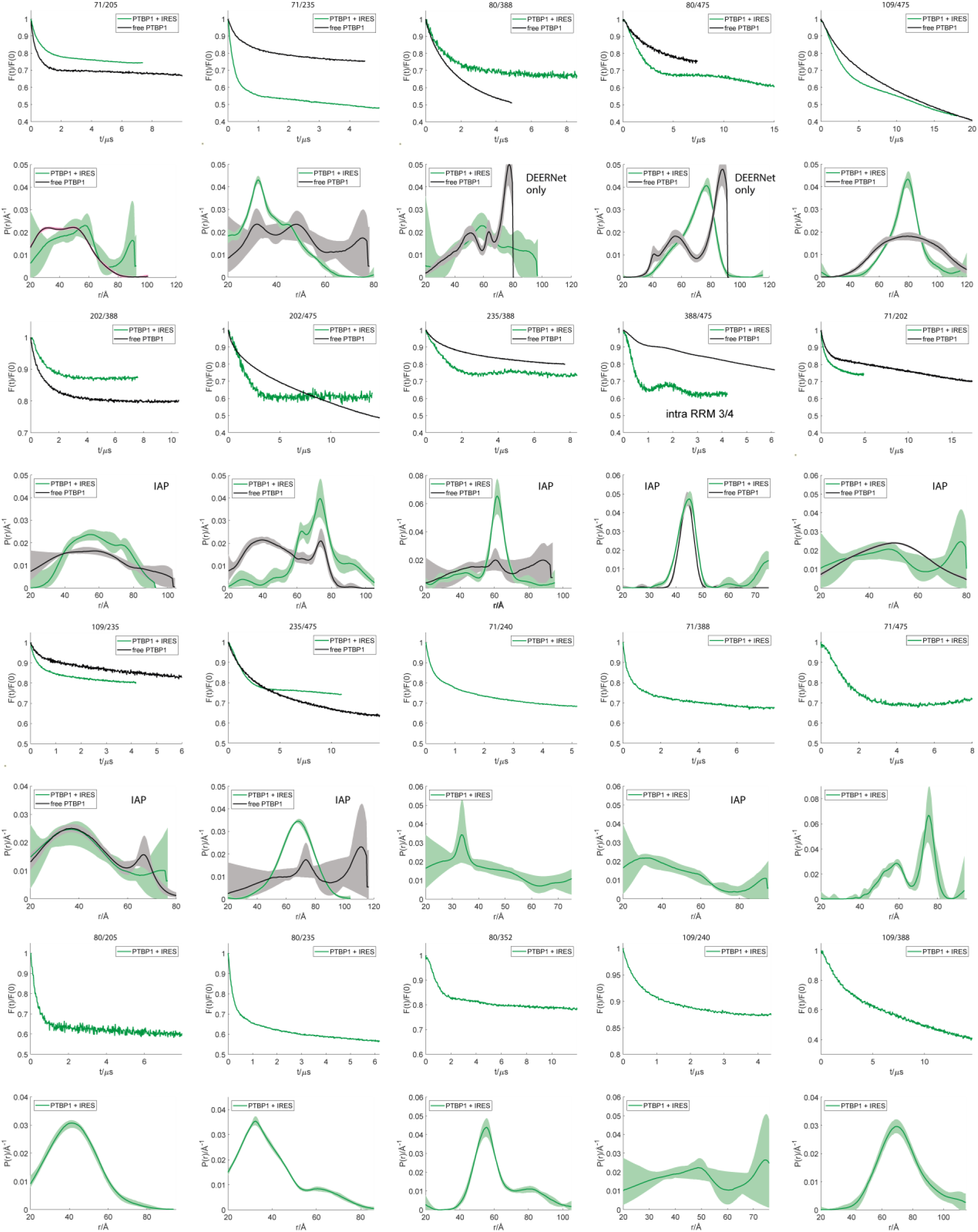
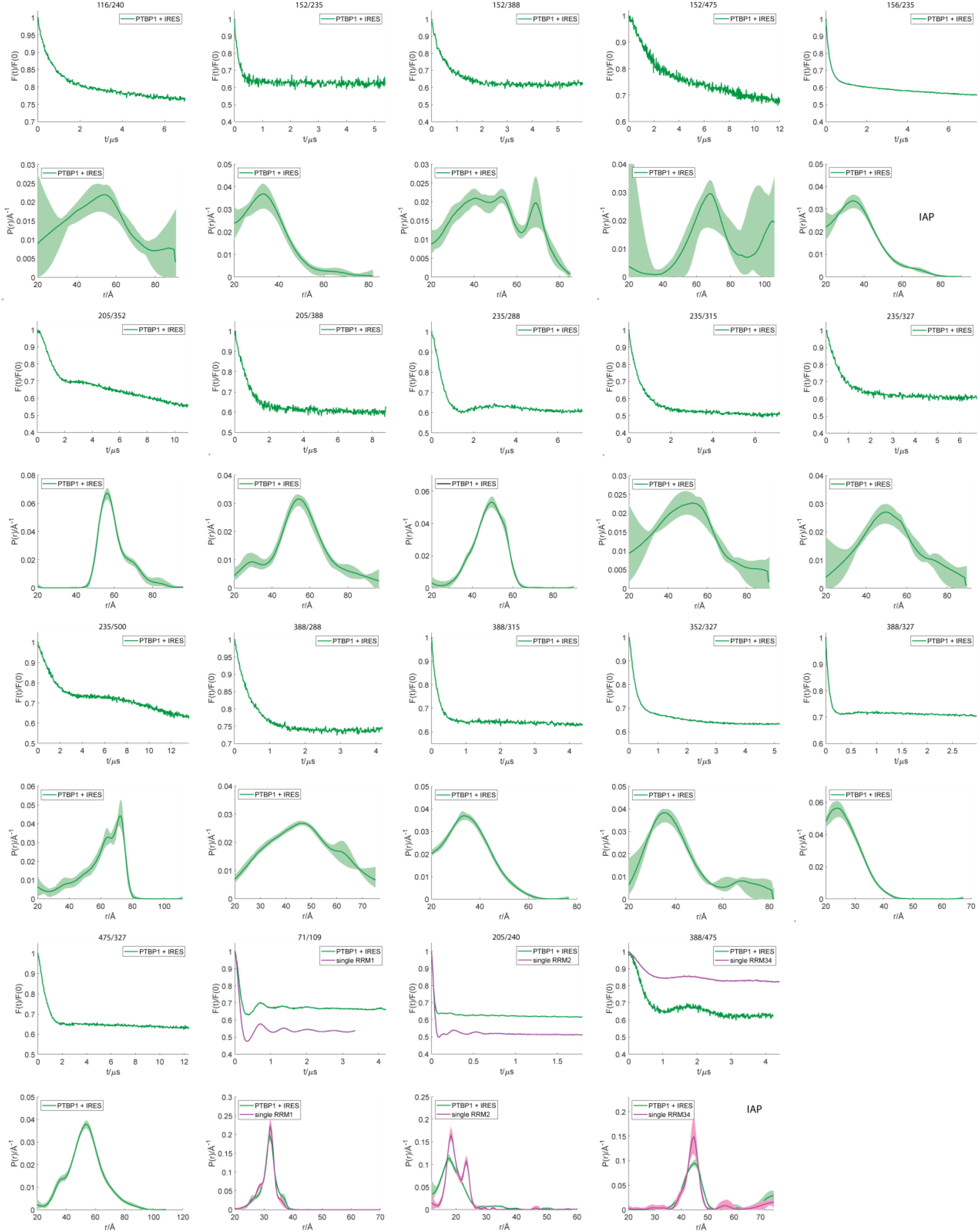
Distance distributions calculated from DEER-EPR data recorded on free PTBP1 and the PTBP1-EMCV complex. Primary time-domain data are shown above the distributions. Green color denotes the complex and black color the free protein. Semi-transparent areas correspond to 95% confidence intervals. Samples with iodoacetamido proxyl spin label are indicated by “IAP”. Distance distributions computed by neural network analysis with DeerNet (86) are indicated.

**Fig. S3.**
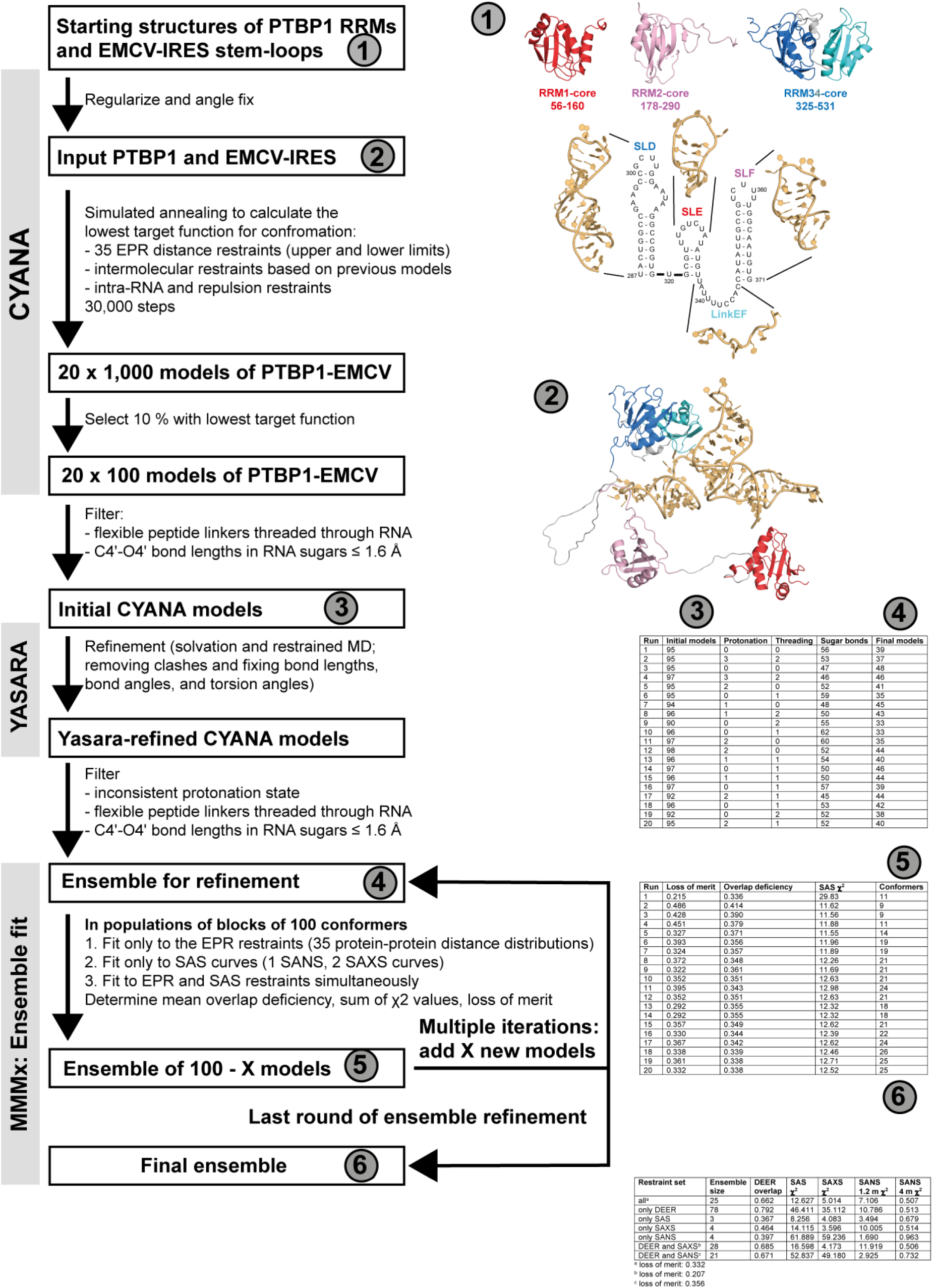
Scheme of the integrative structural modeling approach. For details see Materials and Methods. For ensemble fitting, the number of iterations depends on the number of models in the preliminary ensembles and cannot be predicted.

**Fig. S4.**
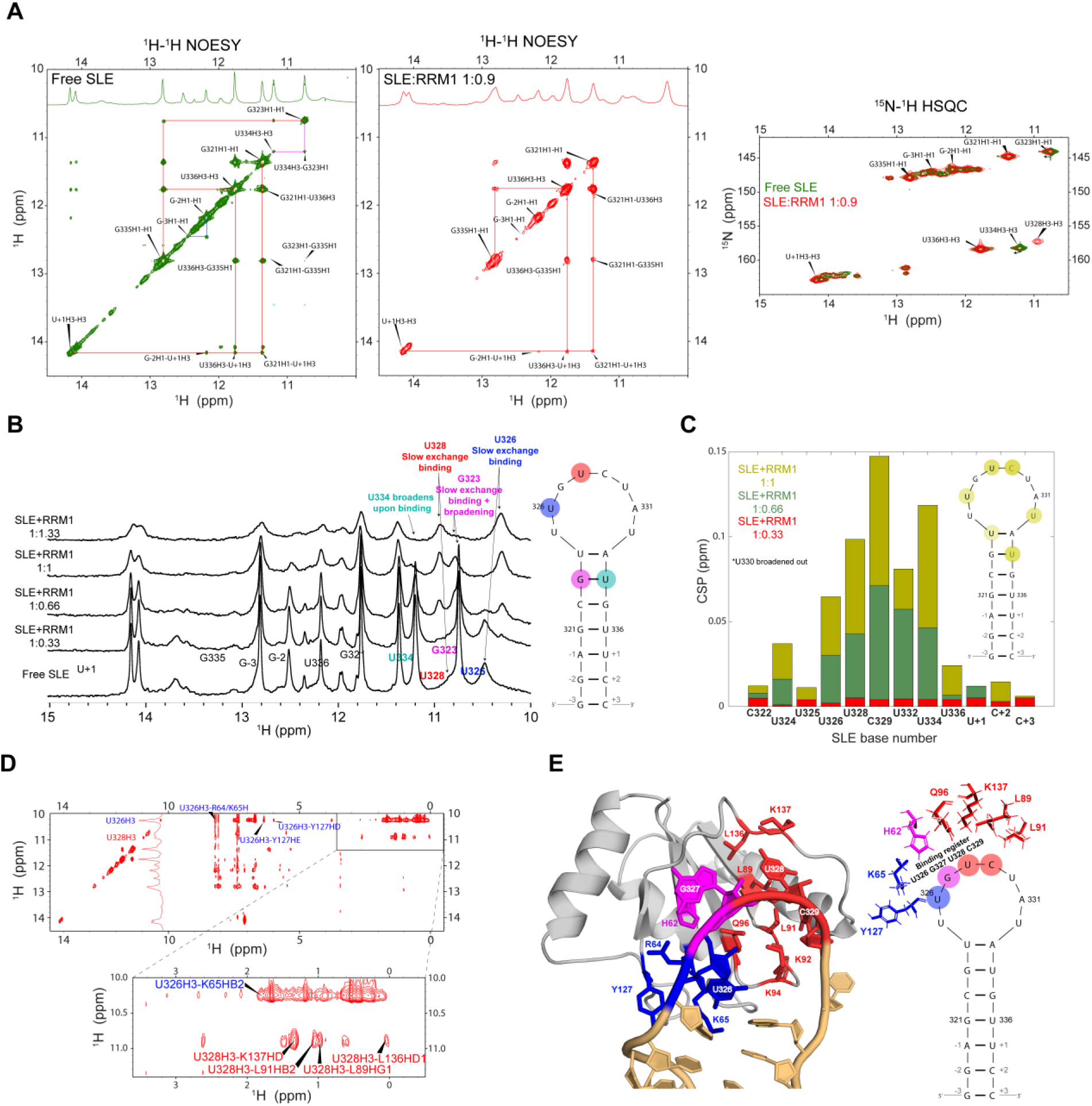
NMR confirmation of the PTBP1 RRM1/SLE binding register. (A) The secondary structure of SLE stem remains unaffected upon binding with RRM1 as shown by 2D ^1^H-^1^H NOESY imino proton sequential walk and 2D ^1^H-^15^ NHSQC spectra at 278 K. Even though upon binding, the G323 and U334 wobble base pair do not give characteristic NOE cross-peak in 2D ^1^H-^1^H NOESY spectra (probably due to signal broadening caused by increased correlation time upon binding), these resonances are still present in a 2D ^1^H-^15^N HSQC spectrum. This indicates that they are still forming a base pair and consequently implies that U324 and A333 also remain base paired despite the absence of an assigned U324-H3 signal. Correlations labeled with asterisk are assigned using a recently developed SMT imino experiment (81). (B) Titration of SLE with RRM1. Perturbation of SLE imino proton signals upon binding at 278 K lead to important conclusions about the binding interface. U326 and U328 are in the slow exchange binding regime and remain sharp due to direct stabilizing interaction with protein. Imino signals of U326 and U328 are assigned according to intermolecular NOE signals with RRM1. G323 and U334 imino proton signals experience additional broadening due to slightly faster chemical exchange of respective imino hydrogens with water. (C) CSP perturbations of H5-H6 correlations. Plot of H5-H6 chemical shift perturbations upon addition of RRM1 calculated from correlations in the ^1^H-^1^H TOCSY experiment. Strongest perturbations are observed for U326, U328, C329, U332 and U334 implying largest conformational changes involving mostly bases within the apical loop upon protein-RNA interaction. (D) Intermolecular L-PROSY NOESY (82), identifying U326 and U328 imino proton signals. Intermolecular NOE contacts involving imino proton resonances acquired using L-PROSY NOESY experiment at 278 K. Cross-peaks detected for the protein can be used to facilitate assignment of two imino proton resonances from the apical loop, U326 and U328. (E) Mapping of binding interface based on NOESY correlations. Multiple intermolecular NOESY experiments acquired in H_2_O and D_2_O (at 278 K and 323 K, respectively) reveal intermolecular correlations between protein and RNA. These intermolecular contacts are highlighted on the 3D structural model. All NMR data support the binding register that is illustrated on the secondary structure of SLE.

**Fig. S5.**
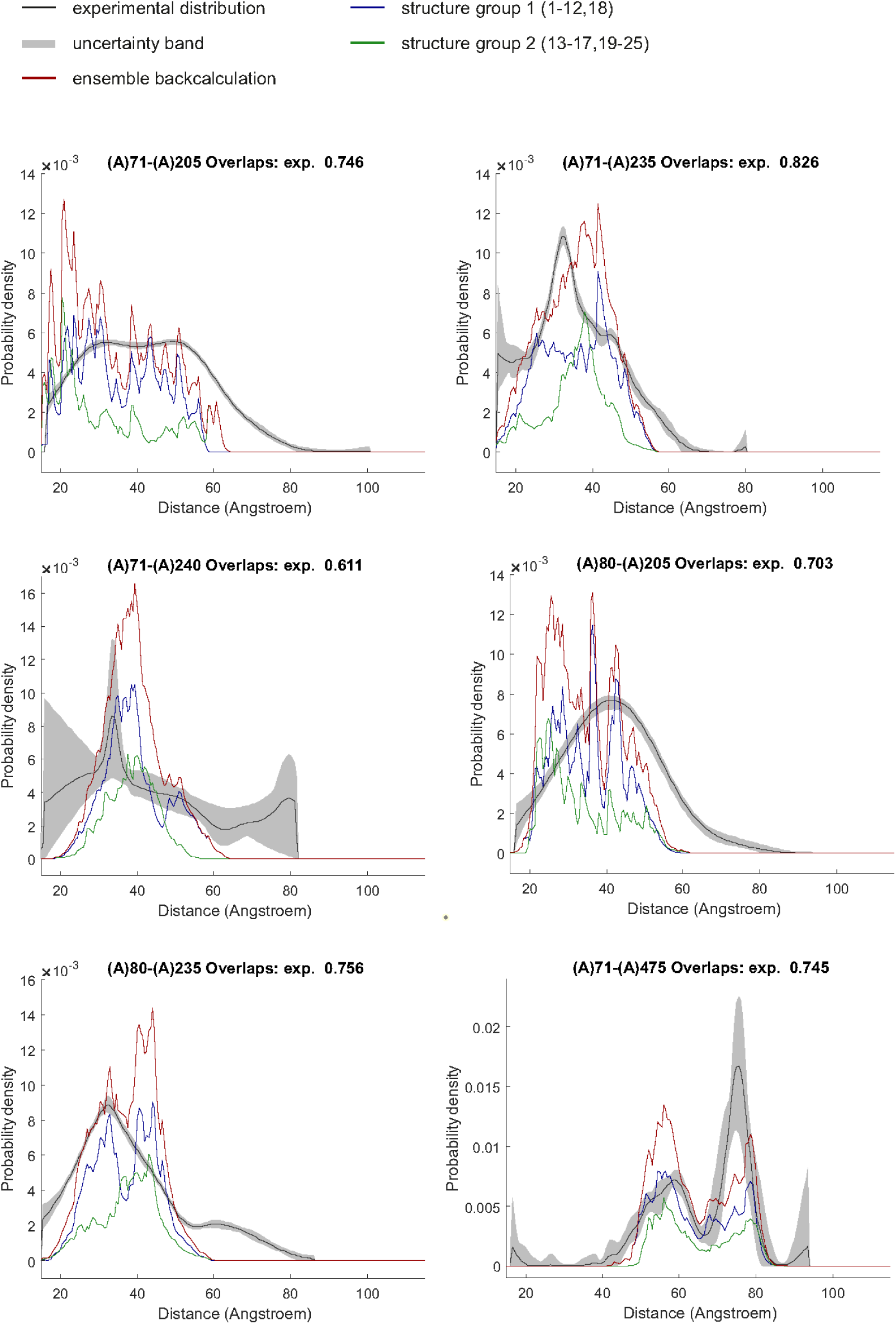
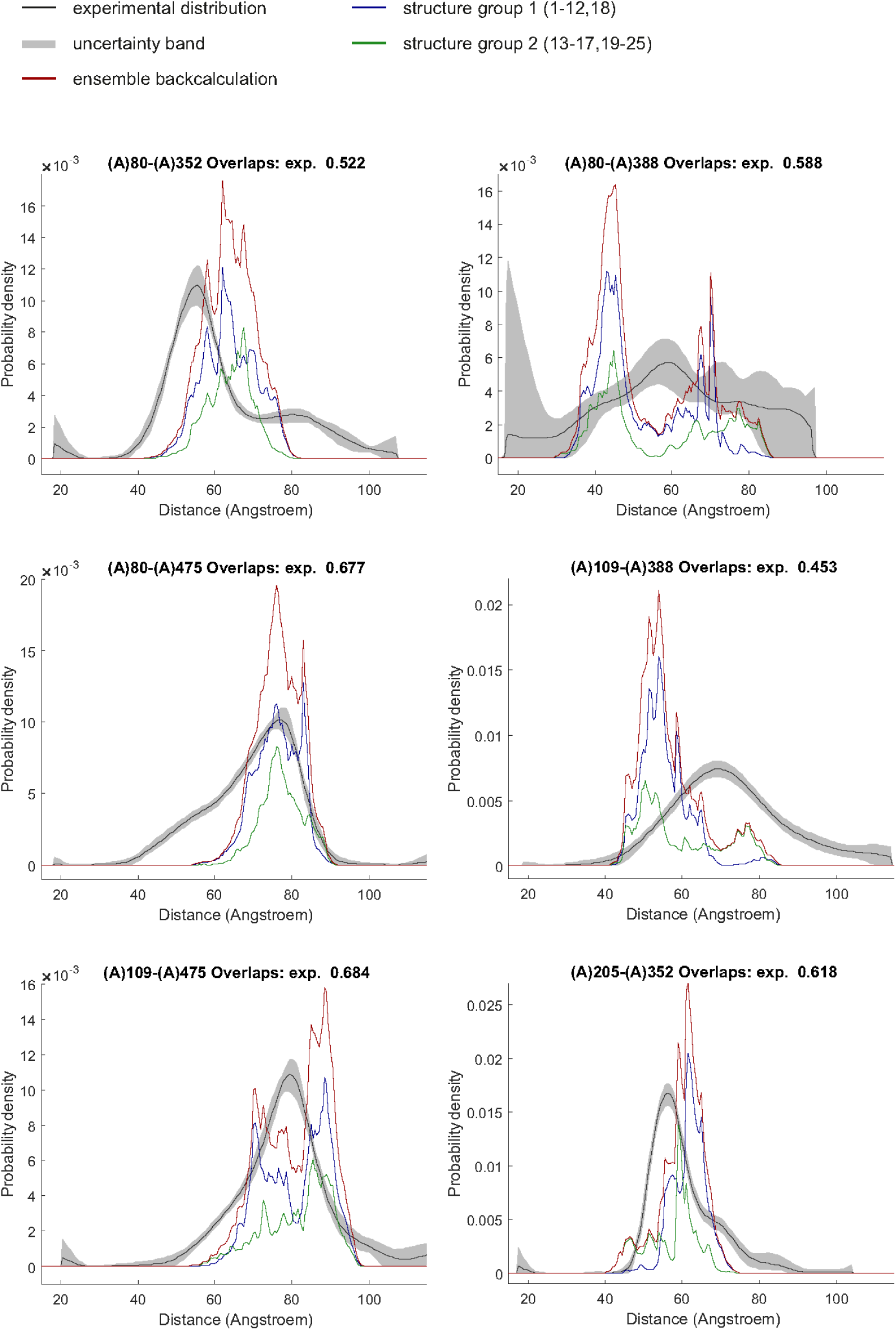
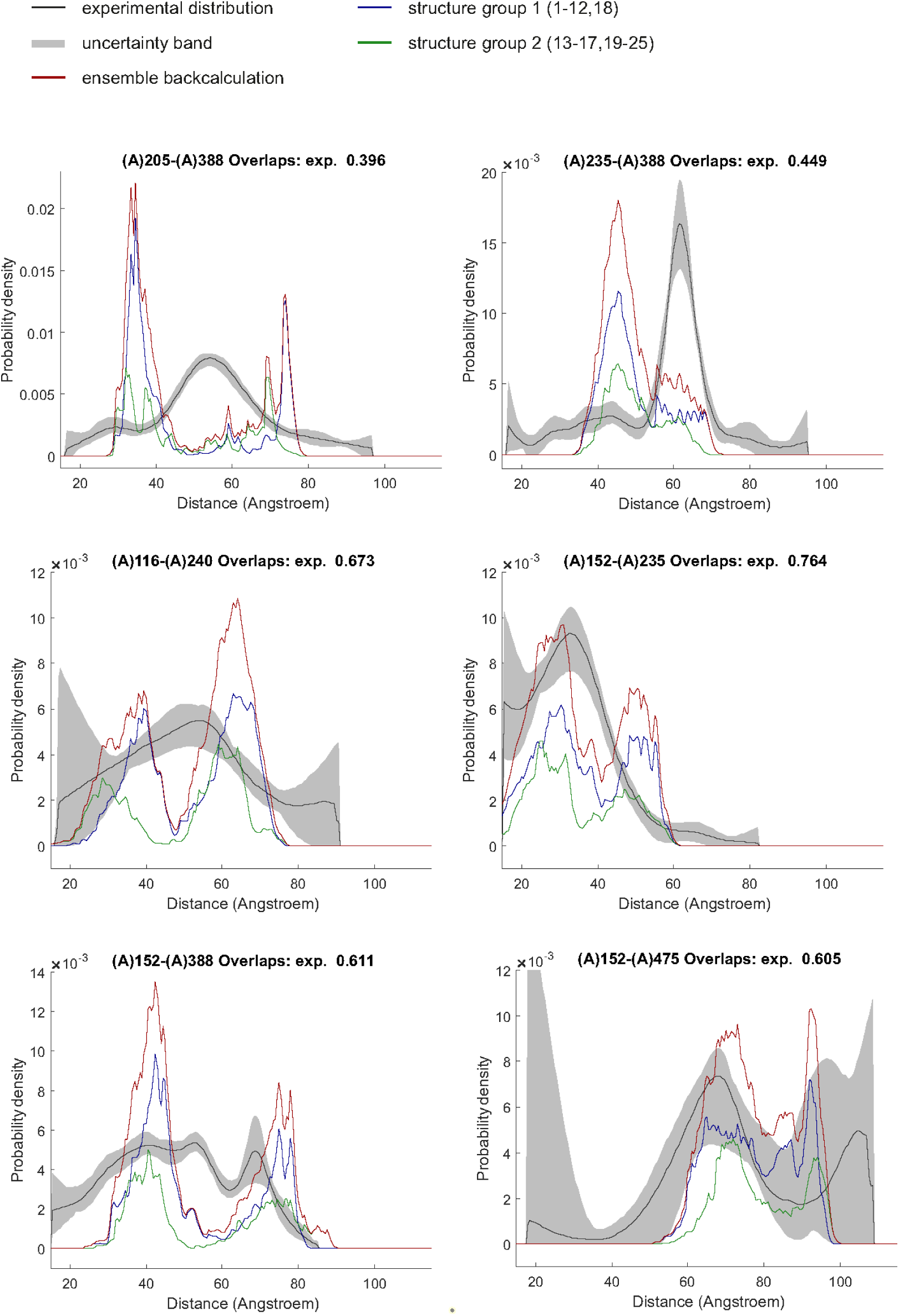
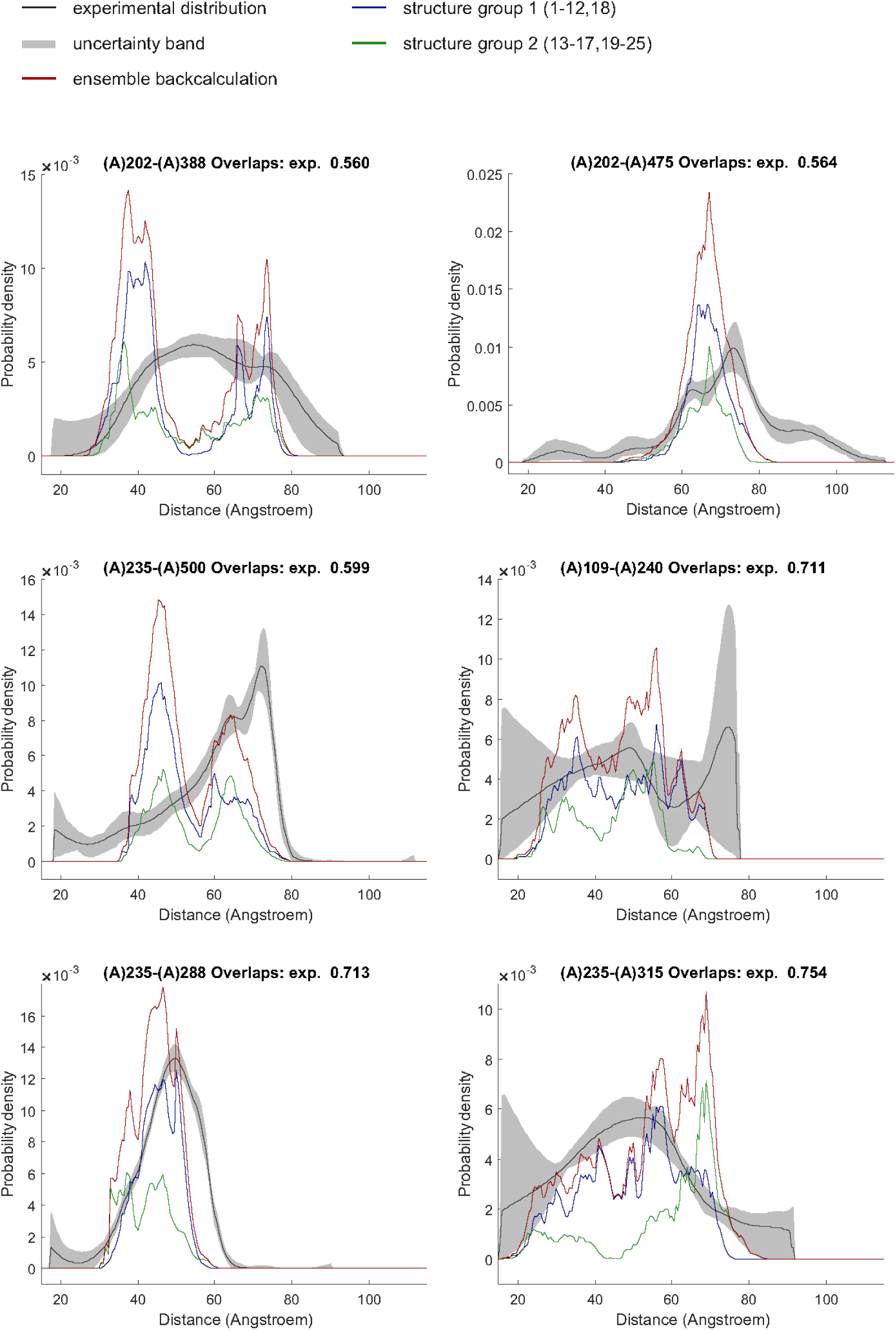
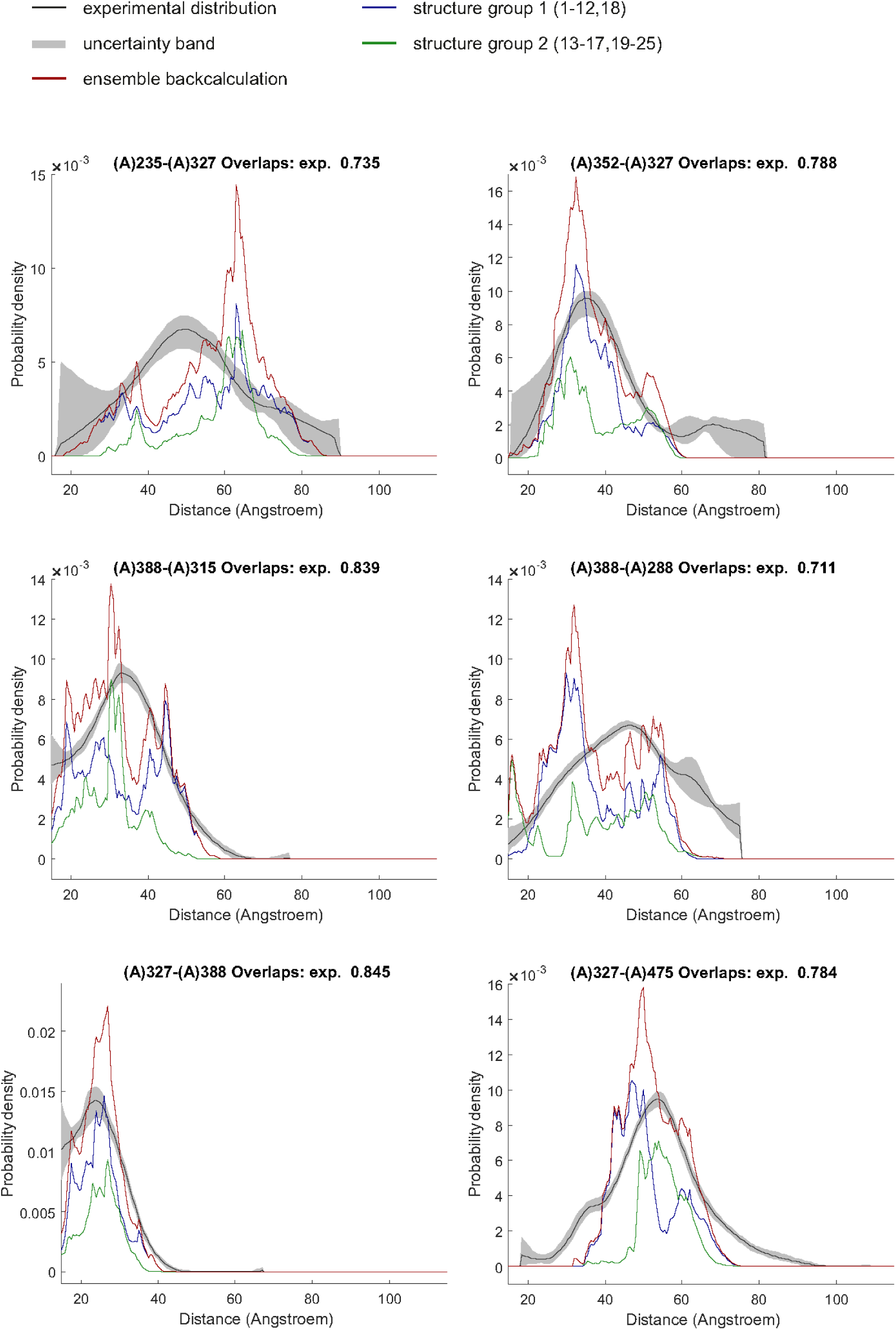
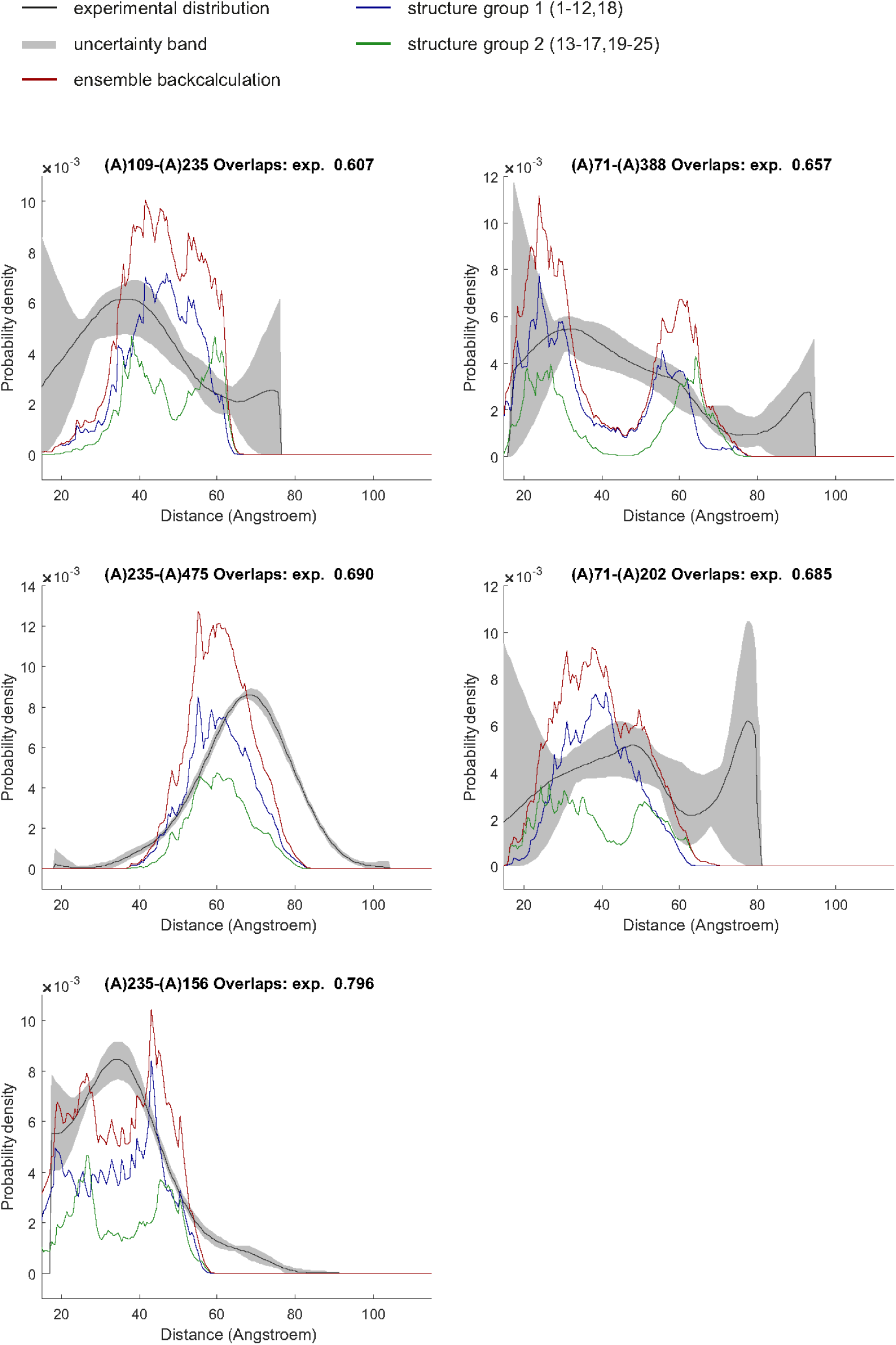
Restraint fit of the integrative PTBP1/EMCV-IRES-DtoF ensemble with DEER distance distributions. Experimentally determined distance distribution (black line) with corresponding uncertainty (grey shading) and predicted for the entire ensemble (red line) or only cis (group 1, purple) and trans (group 2, green) sub-groups, respectively. Overlap values between experimental and predicted distributions for each spin pair are displayed at the top of each panel.

**Fig. S6.**
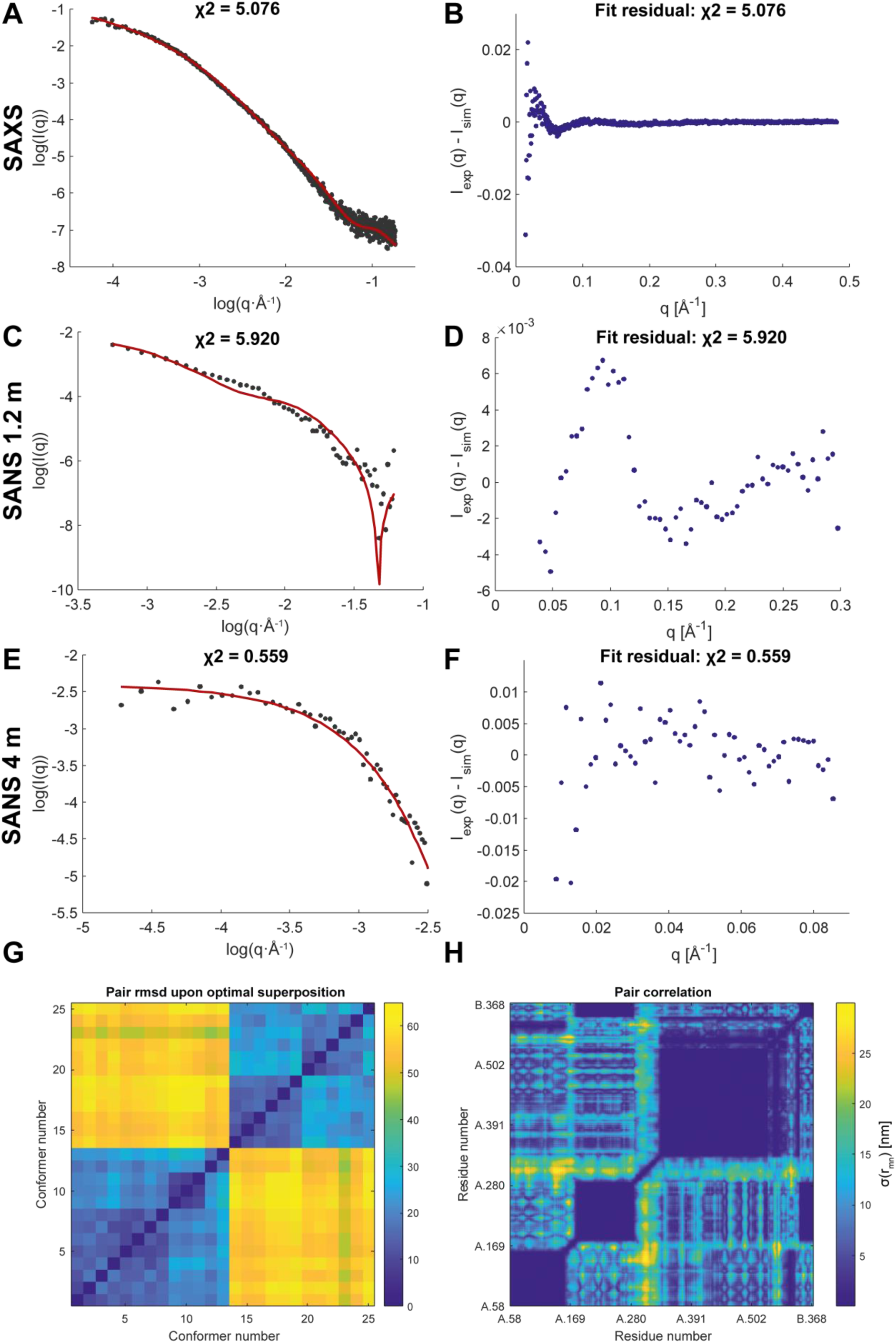
Restraint fit and residuals of the integrative PTBP1/EMCV-IRES-DtoF ensemble with SANS/SAXS data as well as pair-wise correlation. Fit (A) and residual (B) of ensemble with SAXS, SANS 1.2 m (C,D), and SANS 4 m (E,F). Red curve was back-calculated from the ensemble. (G) Conformer pair root mean square deviation (Å). (H) Pair correlation matrix of the ensemble. Distances (Å) were measured between Cα of the two residues.

**Fig. S7.**
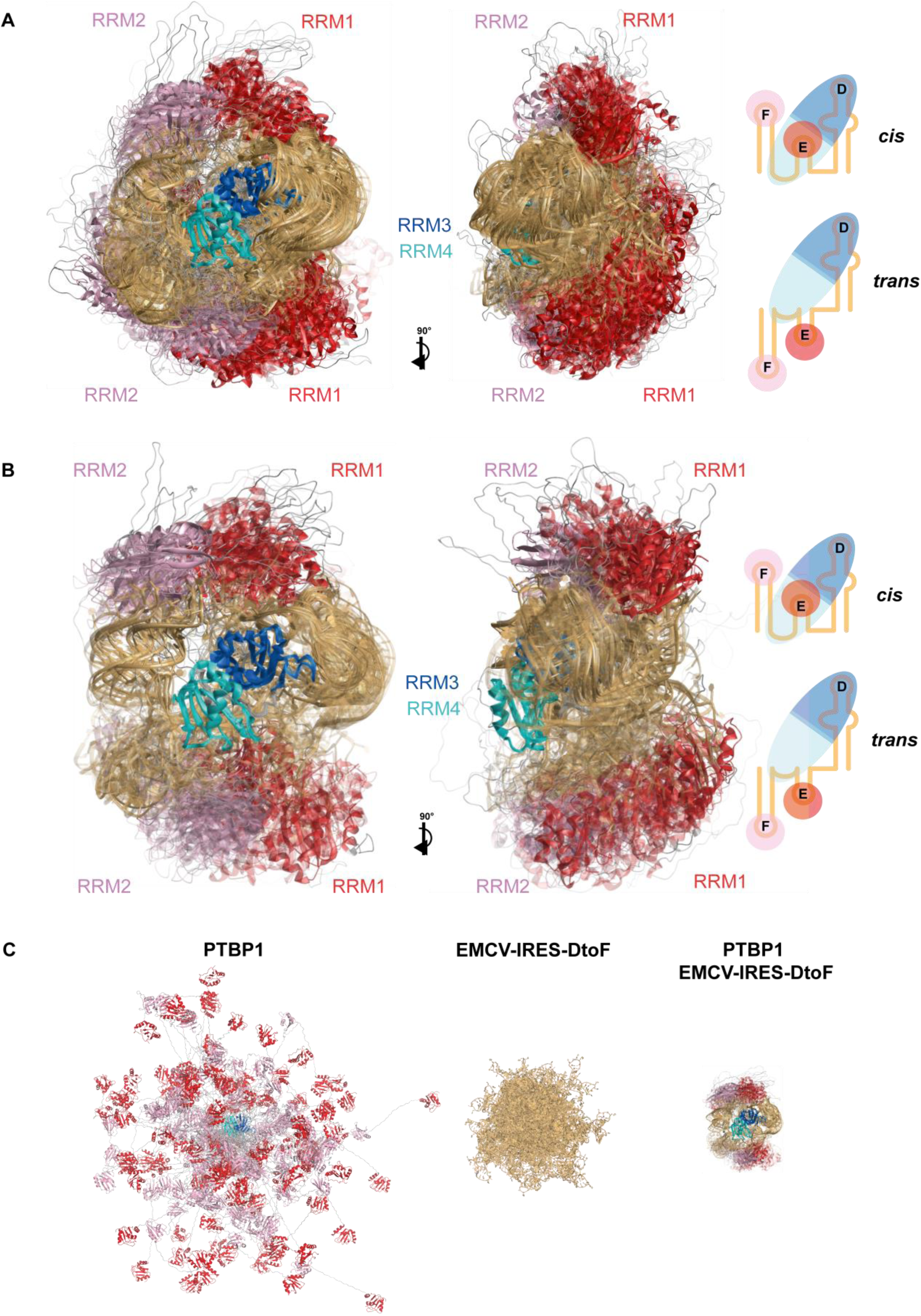
PTBP1/EMCV-IRES-DtoF ensembles obtained with only DEER restraints, after removal of the conformers in the best-fitting ensemble, and unrestrained component ensembles. (A) Structural ensemble of the PTBP1/EMCV-IRES-DtoF complex generated by ensemble fitting only against DEER data. This ensemble has a slightly better fit for all the distance distributions with an overlap between experimental and back-calculated distance distributions between 0.614 and 0.933 with a geometric average of 0.792. Structural ensemble in a conformer population-weighted visualization. The 78 models of the ensemble were superimposed on RRM34. Two views related by a vertical rotation of 90° are shown. As in the integrative structural ensemble, SLE and SLF can be positioned in *cis* or *trans* with respect to SLD, illustrated in schemes on the right. (B) An ensemble obtained in another fit after removing the 25 conformers that were included in the best-fitted integrative ensemble. This ensemble has 30 conformers. Fit of the small-angle scattering curves has deteriorated (χ^2^: 15.542) and fit of the DEER distance distributions (geometric average: 0.690) and loss of merit (0.266) have even slightly improved. (C) Unrestrained PTBP1 and EMCV-IRES-DtoF ensembles (100 conformers) superimposed on RRM34 or SLF, respectively, in comparison to the integrative ensemble of the PTBP1/EMCV-IRES-DtoF complex.

**Fig. S8.**
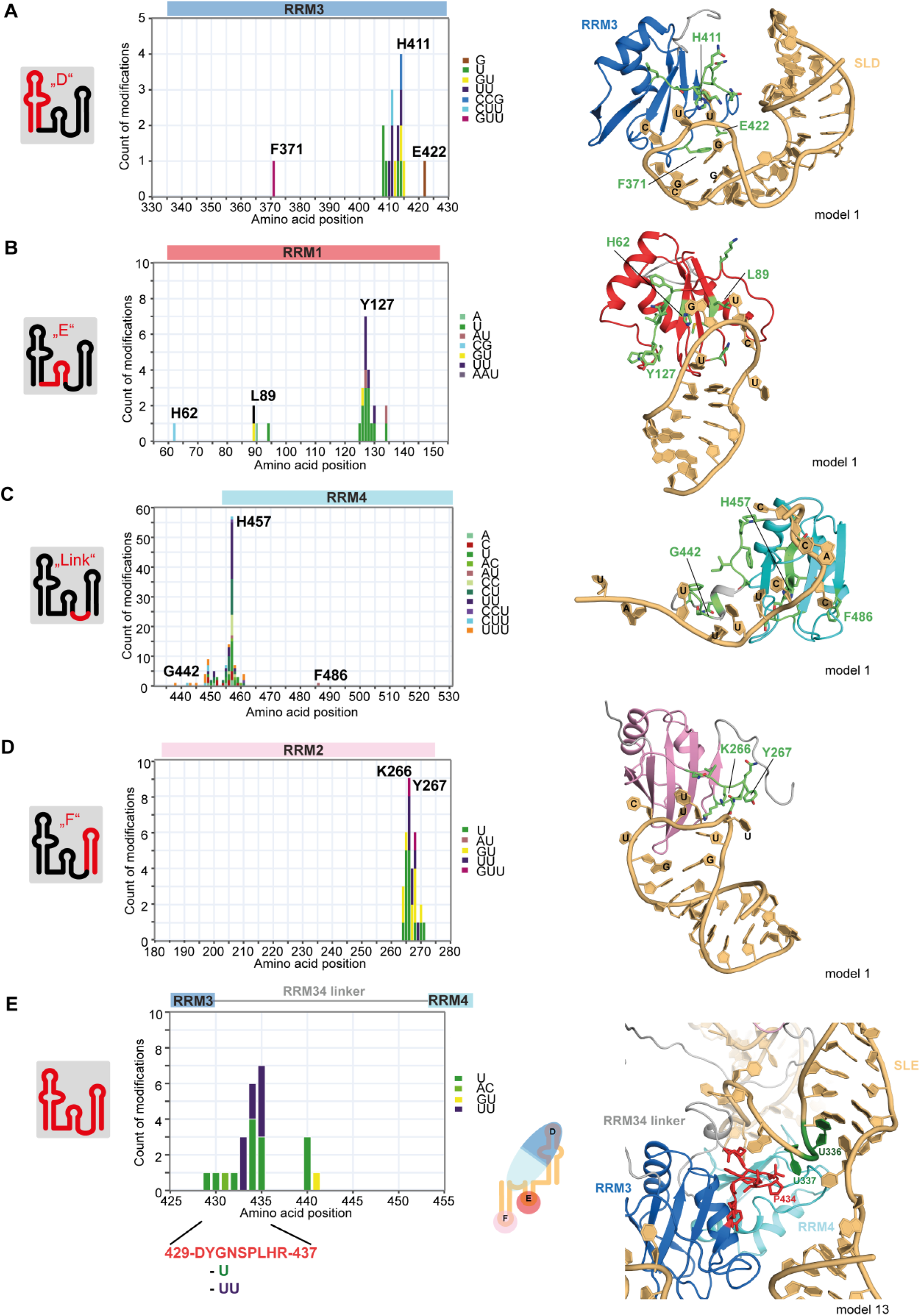
Comparison of structural ensemble with updated CLIR-MS/MS data of PTBP1 in complex with segmentally or uniformly isotope labeled RNA. CLIR-MS/MS data were re-analyzed with an updated analysis pipeline. (39, 76). (A)-(D) Cross-link identification detected at each labeling site (D, E, Link, F) for the respective RRM plotted on the sequence of the protein. Colors indicate type of cross-link modification (left). Cross-links shown on the structural models of the corresponding sub-complexes (right). (E) Cross-links identified in the linker between RRM3 and RRM4.

**Fig. S9.**
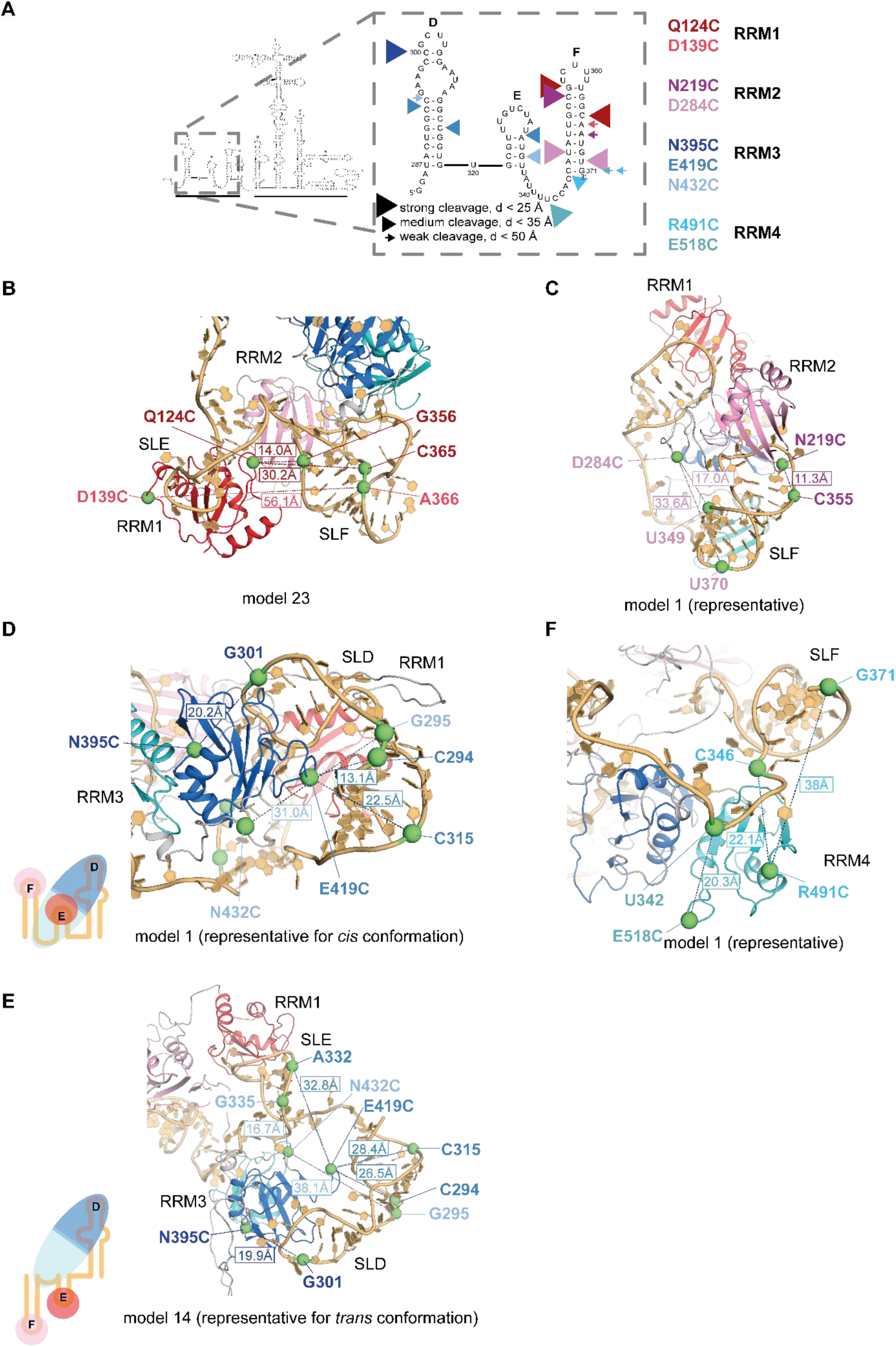
Comparison of structural ensemble with hydroxyl radical probing data. (A) Strong, medium and weak cleavages detected for PTBP1 in complex with EMCV-IRES (43). (B) RRM1 cleavages on SLF are generally not consistent with the mapping of the domain on SLE, however, in a few models (e.g. model 23) residues are close enough to induce a cleavage in SLF. (C) RRM2 cleavages in SLD fit well with the mapping of the domain to this stem-loop and in all the models, the distances are consistent with hydroxyl radical cleavage data. In contrast to previous assumptions, RRM2 does not bind the stem of SLF. (D) RRM3 cleavages on SLD are fully consistent with the mode of binding to SLD in all the models, some cleavages on SLE are rationalized by close distances to SLE in the *trans* conformation (E). RRM4 cleavages to the LinkEF sequence and the basis of SLF fit well with all models in the ensemble.

**Fig. S10.**
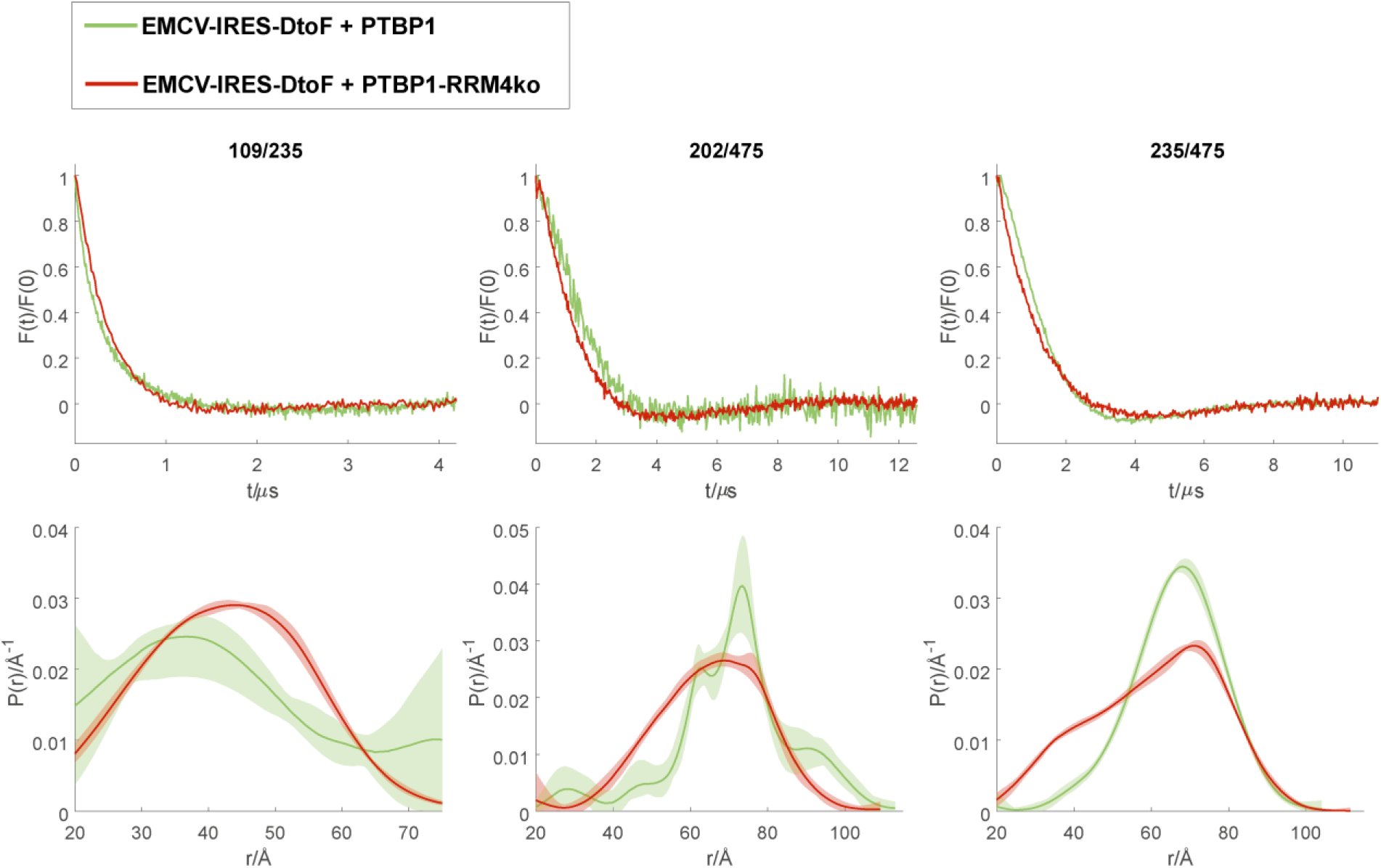
DEER distance measurements of RRM4ko mutants. DEER measurement of WT PTBP1 and construct with mutated binding interface of RRM4 (RRM4ko, red) in complex with EMCV-IRES-DtoF. Three spin-pairs reflecting RRM1-RRM2 (109/235) and RRM2-RRM34 (202/475 and 235/475) distances, respectively. Top panel, form factor and lower panel, distance distribution.

**Table S1.**
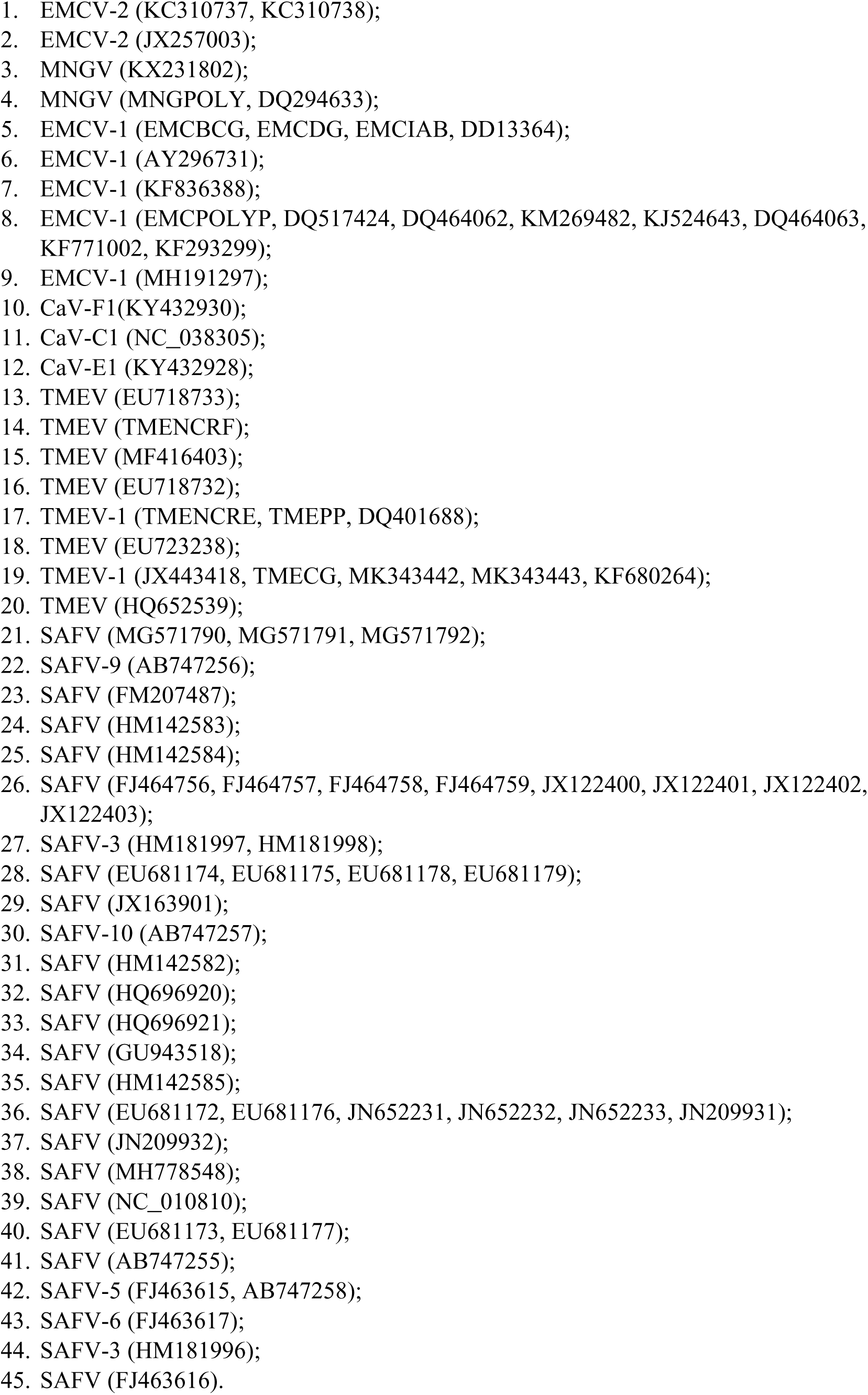
List of cardiovirus sequences used in the alignment and their accession numbers.

**Table S2.**
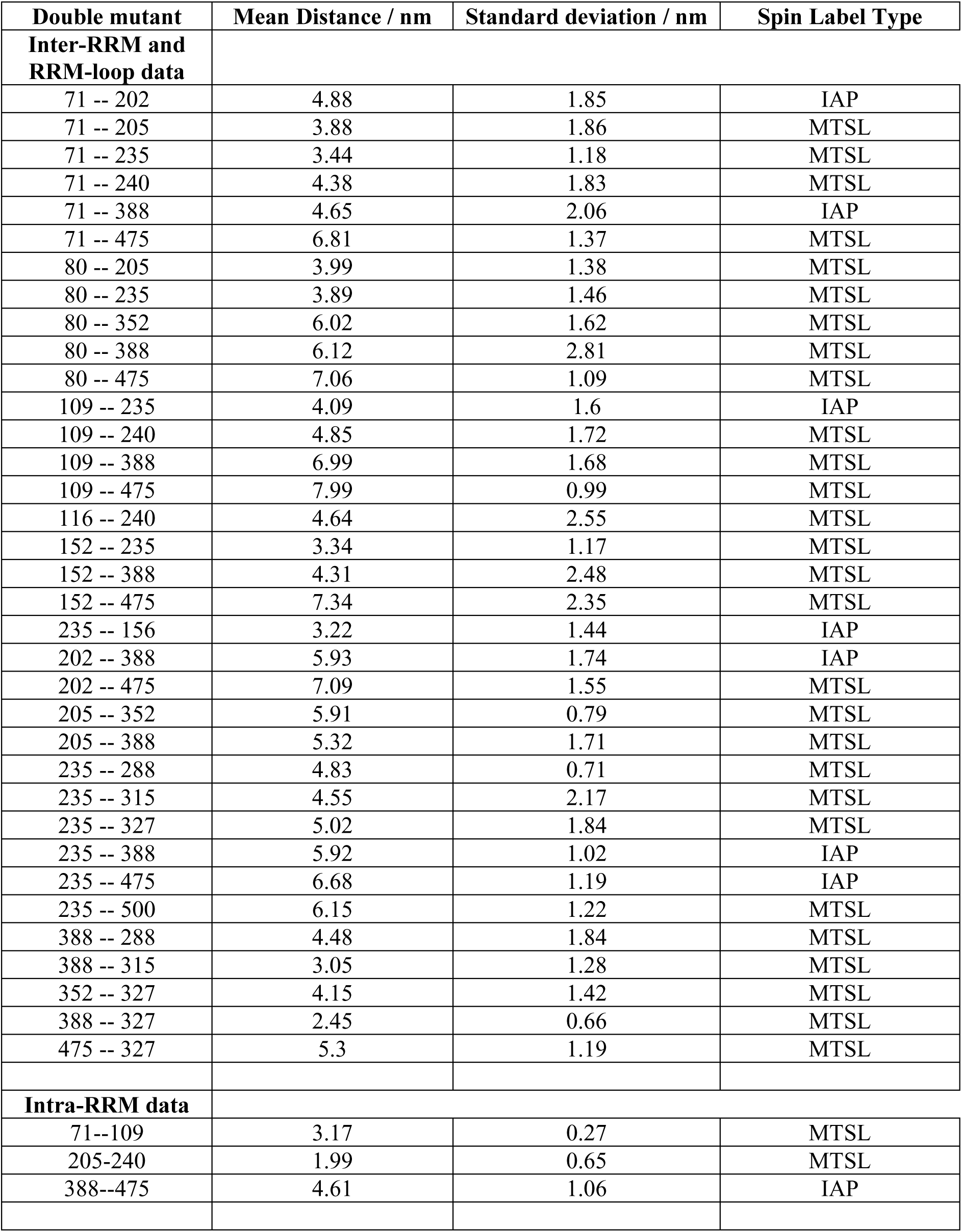
Distance distribution measurements of doubly-labeled PTBP1 in complex with EMCV-IRES-DtoF.

| Double mutant | Mean Distance / nm | Standard deviation / nm | Spin Label Type |
| --- | --- | --- | --- |
| **Inter-RRM and RRM-loop data** |  |  |  |
| 71 -- 202 | 4.88 | 1.85 | IAP |
| 71 -- 205 | 3.88 | 1.86 | MTSL |
| 71 -- 235 | 3.44 | 1.18 | MTSL |
| 71 -- 240 | 4.38 | 1.83 | MTSL |
| 71 -- 388 | 4.65 | 2.06 | IAP |
| 71 -- 475 | 6.81 | 1.37 | MTSL |
| 80 -- 205 | 3.99 | 1.38 | MTSL |
| 80 -- 235 | 3.89 | 1.46 | MTSL |
| 80 -- 352 | 6.02 | 1.62 | MTSL |
| 80 -- 388 | 6.12 | 2.81 | MTSL |
| 80 -- 475 | 7.06 | 1.09 | MTSL |
| 109 -- 235 | 4.09 | 1.6 | IAP |
| 109 -- 240 | 4.85 | 1.72 | MTSL |
| 109 -- 388 | 6.99 | 1.68 | MTSL |
| 109 -- 475 | 7.99 | 0.99 | MTSL |
| 116 -- 240 | 4.64 | 2.55 | MTSL |
| 152 -- 235 | 3.34 | 1.17 | MTSL |
| 152 -- 388 | 4.31 | 2.48 | MTSL |
| 152 -- 475 | 7.34 | 2.35 | MTSL |
| 235 -- 156 | 3.22 | 1.44 | IAP |
| 202 -- 388 | 5.93 | 1.74 | IAP |
| 202 -- 475 | 7.09 | 1.55 | MTSL |
| 205 -- 352 | 5.91 | 0.79 | MTSL |
| 205 -- 388 | 5.32 | 1.71 | MTSL |
| 235 -- 288 | 4.83 | 0.71 | MTSL |
| 235 -- 315 | 4.55 | 2.17 | MTSL |
| 235 -- 327 | 5.02 | 1.84 | MTSL |
| 235 -- 388 | 5.92 | 1.02 | IAP |
| 235 -- 475 | 6.68 | 1.19 | IAP |
| 235 -- 500 | 6.15 | 1.22 | MTSL |
| 388 -- 288 | 4.48 | 1.84 | MTSL |
| 388 -- 315 | 3.05 | 1.28 | MTSL |
| 352 -- 327 | 4.15 | 1.42 | MTSL |
| 388 -- 327 | 2.45 | 0.66 | MTSL |
| 475 -- 327 | 5.3 | 1.19 | MTSL |
| **Intra-RRM data** |  |  |  |
| 71--109 | 3.17 | 0.27 | MTSL |
| 205-240 | 1.99 | 0.65 | MTSL |
| 388--475 | 4.61 | 1.06 | IAP |

